# Aldosterone drives HSD2 neuron activity and molecular remodeling of synaptic function to promote sodium appetite

**DOI:** 10.64898/2026.10.21.759848

**Authors:** Miriam C McDonough, Ahmet Kuralay, Trevor B Butler, Alysia B Berns, Madelin R Schwager, Silvia Gasparini, Joel C Geerling, Jon M Resch

## Abstract

Sodium deficiency increases aldosterone, which signals through the mineralocorticoid receptor (MR) to restrict sodium excretion and promote salt appetite. Despite aldosterone’s importance in maintaining hydromineral balance, its role in regulating neuronal function is poorly understood. Here, we define the transcriptional response to aldosterone in neurons and demonstrate that MR signaling in HSD2 neurons is essential for aldosterone-induced sodium appetite. Disruption of MR in HSD2 neurons impaired sodium appetite following deficiency and abolished saline intake in response to aldosterone. Single-nucleus RNA sequencing of the caudal nucleus of the solitary tract (NTS) revealed that aldosterone selectively regulates gene expression in HSD2 neurons, with enrichment for transcripts associated with synaptic function. Consistent with this, aldosterone increased the paired-pulse ratio at HSD2 neuron synapses, indicating reduced synaptic depression during repeated stimulation. Notably, *Sv2c*, encoding synaptic vesicle glycoprotein 2C, exhibited the greatest increase in expression after aldosterone treatment, and its disruption in HSD2 neurons reduced saline ingestion, identifying SV2C as a mediator of mineralocorticoid-induced sodium appetite. Together, these findings reveal a selective aldosterone-induced transcriptional program associated with enhanced synaptic transmission in HSD2 neurons and identify a molecular mechanism through which aldosterone promotes sodium appetite.

## Introduction

Sodium is the primary determinant of extracellular fluid volume and must therefore be tightly controlled. A key regulator of sodium balance is aldosterone, which is released from the adrenal cortex in response to sodium deficiency. Aldosterone signals through the mineralocorticoid receptor (MR) to facilitate sodium retention^1^ and promote sodium ingestion^2–6^. Accordingly, aldosterone infusion into the brain increases sodium intake, while interference with MR signaling diminishes it^5,7,8^. Aldosterone sensitivity requires expression of both MR and the glucocorticoid-inactivating enzyme 11β-hydroxysteroid dehydrogenase type II (HSD2)^9–11^, which in the brain is uniquely expressed by a population of neurons in the nucleus of the solitary tract (NTS)^12,13^. HSD2 neurons are activated by mineralocorticoids and drive sodium intake^3,6,14–16^, whereas their ablation abolishes aldosterone-mediated sodium ingestion^3^. Thus, aldosterone may act primarily through HSD2 neurons to drive sodium appetite.

Despite the effects of mineralocorticoids on appetite and mood-related behaviors^2,8,17–26^, neuronal aldosterone-MR signaling has received little mechanistic investigation. Chronic aldosterone infusion in mice induces cell-autonomous action potential firing of HSD2 neurons in *ex vivo* brain slice recordings^14^, suggesting that aldosterone-MR signaling promotes intrinsic activity via genomic mechanisms. Single-cell RNA sequencing (scRNA-seq) of HSD2 neurons following sodium deprivation identified widespread gene expression changes, but these cannot be directly attributed to aldosterone, and few of the differentially expressed genes (DEGs) have been established as MR-regulated^14^. Moreover, the transcriptional effects of MR signaling vary across cell types^27–29^, making it difficult to extrapolate mechanisms identified in other aldosterone-sensitive tissues to neurons. This leaves the genomic response to aldosterone in neurons unresolved.

The unique aldosterone sensitivity of HSD2 neurons provides an opportunity to identify the molecular mechanisms through which aldosterone drives salt appetite and offer insight into the transcriptional program coordinated by MR signaling in neurons. In this study, we utilize genetic approaches to establish the importance of aldosterone-MR signaling in HSD2 neurons for sodium appetite. Using HSD2 neuron-enriched single-nucleus RNA sequencing (snRNA-seq) of the caudal NTS, we identify a highly selective transcriptional response to aldosterone, with enrichment for genes associated with synaptic function. Guided by our transcriptomic analysis, we demonstrate that aldosterone alters synaptic transmission from HSD2 neurons and identify synaptic vesicle glycoprotein 2C (SV2C) as an MR-regulated mediator of HSD2 neuron function and sodium appetite.

## Results

### MR deletion from VGLUT2^+^ neurons eliminates HSD2 expression and impairs sodium appetite

HSD2 neurons exhibit strong enrichment for *Nr3c2*, which encodes MR, and chronic aldosterone infusion induces sodium intake and c-Fos expression in HSD2 neurons (**Figures 1A-1C**)^3,6,12,14,16,30^. Despite the established role of HSD2 neurons in sodium appetite, how loss of MR signaling affects their function and sodium intake has not been examined. Conditional deletion of *Nr3c2* from all HSD2-expressing cells would disrupt MR signaling in the kidney and cause severe salt wasting^31,32^. Therefore, we generated nervous system-specific MR knockouts by crossing *Nr3c2*^lox/lox^ mice^33^ to either *Nestin*-Cre^34^ or *Syn1*-Cre^35^ strains. Both lines successfully eliminated robust *Nr3c2* expression in the hippocampus (**Figures S1A and S1D**). However, *Nestin*-Cre::*Nr3c2*^lox/lox^ mice retained *Nr3c2* expression in multiple brain regions, often in cholinergic motor neurons (**Figures S1B and S1E**). More importantly, both conditional knockout lines exhibited incomplete deletion of *Nr3c2* from HSD2 neurons, precluding their use to determine whether MR is required in these neurons for sodium appetite (**Figures S1C and S1F**).

**Figure 1:**
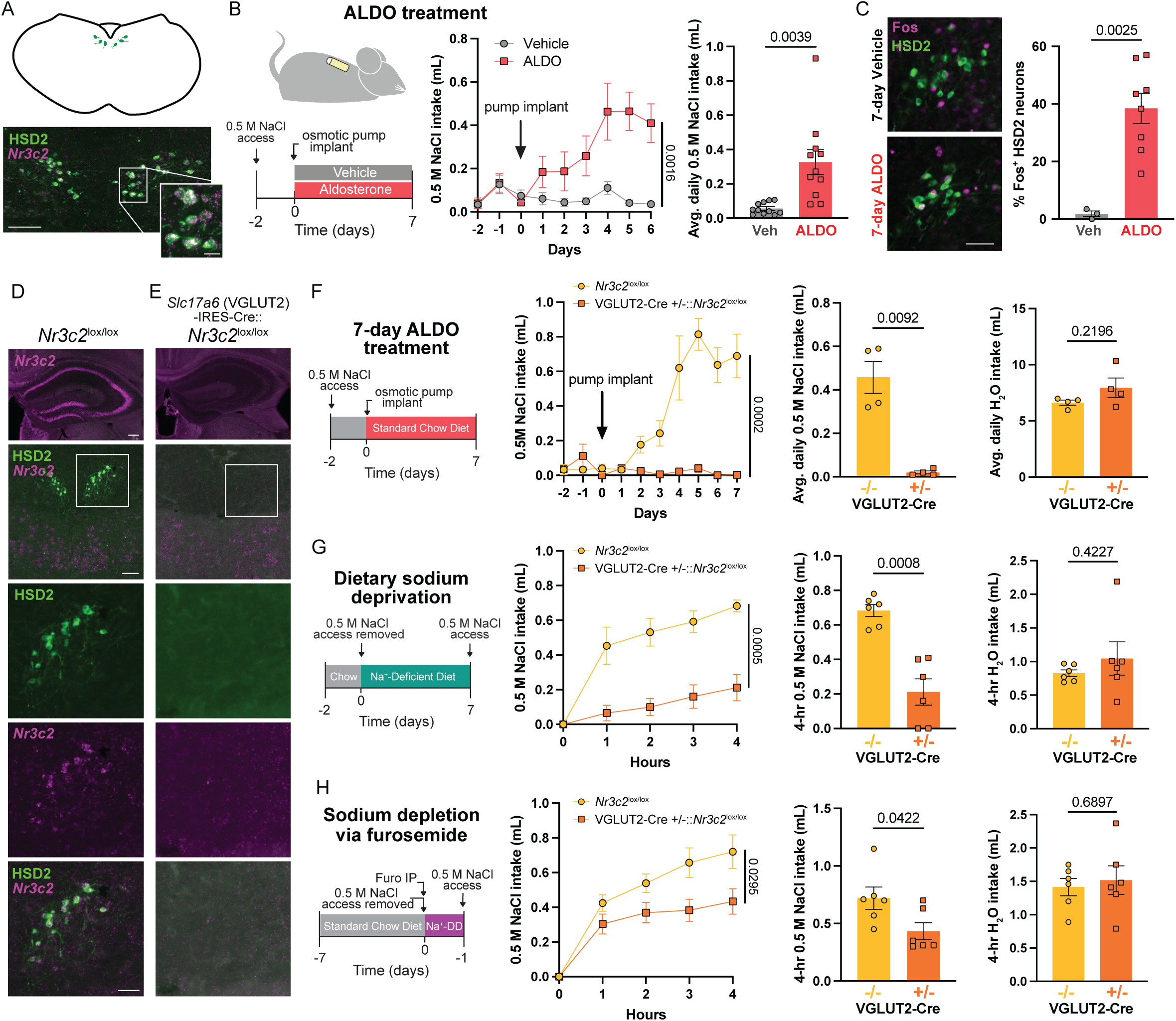
Developmental deletion of *Nr3c2* from VGLUT2^+^ neurons eliminates HSD2 expression in the NTS and diminishes sodium appetite. **A)** Schematic showing HSD2 location in the hindbrain (top) and representative histology showing colocalization of HSD2 immunofluorescence (IF) and *Nr3c2* fluorescence *in situ* hybridization (FISH) in the NTS (bottom; scale bar = 100 µm). Panel inset shows higher magnification image (scale bar = 20 µm). **B)** Schematic describing the aldosterone-induced sodium appetite model (left), 0.5 M NaCl intake in mice during 6 days of vehicle or aldosterone administration (middle) and average 0.5 M NaCl intake per day after aldosterone treatment (right; n = 11 mice/group). **C)** Representative images showing c-Fos expression in HSD2 neurons from 7-day vehicle-treated (n = 3; top left) or aldosterone-treated (n = 8; bottom left) mice and the percentage of HSD2 neurons expressing c-Fos (right; scale bar = 50 μm). **D-E)** Representative brains sections from *Nr3c2*^lox/lox^ **(D)** and VGLUT2-IRES-Cre::*Nr3c2*^lox/lox^ **(E)** mice showing, in descending order: *Nr3c2* in the hippocampus (scale bar = 200 μm); low-magnification view of the dorsal vagal complex with HSD2 IF and *Nr3c2* FISH labeling (scale bar = 100 μm); high-magnification view of HSD2 IF in boxed regions, high-magnification view of *Nr3c2* FISH in boxed regions, and high-magnification view of both HSD2 IF and *Nr3c2* FISH (scale bar = 50 μm). **F)** Showing, left to right: schematic describing the aldosterone-induced sodium appetite model, 0.5 M NaCl intake in *Nr3c2*^lox/lox^ and VGLUT2-IRES-Cre::*Nr3c2*^lox/lox^ mice during 7 days of aldosterone treatment, daily average of 0.5 M NaCl intake, and daily average of H_2_O intake (n = 4 mice/group). ALDO = aldosterone. **G)** Showing, left to right: schematic describing the dietary sodium deprivation sodium appetite model, 0.5 M NaCl intake in *Nr3c2*^lox/lox^ and VGLUT2-IRES-Cre::*Nr3c2*^lox/lox^ mice over 4 hours following sodium deprivation, total 0.5 M NaCl intake in 4 hours, and total H_2_O intake in 4 hours (n = 6 mice/group). **H)** Showing, left to right: schematic describing the furosemide-induced sodium depletion sodium appetite model, 0.5 M NaCl intake in *Nr3c2*^lox/lox^ and VGLUT2-IRES-Cre::*Nr3c2*^lox/lox^ mice over 4 hours following sodium depletion via furosemide, total 0.5 M NaCl intake in 4 hours, and total H_2_O intake in 4 hours (n = 6 mice/group). Furo = furosemide; Na^+^-DD = Na^+^-deficient diet. All data are represented as mean ± SEM. Statistical analysis of saline intake over time between groups was performed using a two-way repeated-measures ANOVA with Geisser-Greenhouse correction. All other statistical analyses were performed with a Welch’s t-test. p < 0.05 was considered statistically significant.

As an alternative, we conditionally deleted *Nr3c2* from neurons expressing the vesicular glutamate transporter 2 (*Slc17a6;* VGLUT2), which include HSD2 neurons^12,14^. VGLUT2-IRES-Cre::*Nr3c2*^lox/lox^ mice again showed loss of *Nr3c2* expression in the hippocampus, but crucially also lacked prominent *Nr3c2* expression in the NTS, consistent with successful deletion of *Nr3c2* from VGLUT2^+^ neurons. Unexpectedly, however, HSD2 immunoreactivity was completely absent in the NTS of these mice (**Figures 1D and 1E**). These results suggest that MR is required for HSD2 expression or, alternatively, for the development or survival of HSD2 neurons.

Although HSD2 neuron ablation impairs sodium appetite^3,14,15^, it is unknown whether developmental MR deletion and the accompanying loss of HSD2 expression produce a similar phenotype. We therefore examined sodium intake in VGLUT2-IRES-Cre::*Nr3c2*^lox/lox^ mice using three standard approaches. First, we measured sodium appetite driven by subcutaneous administration of aldosterone for seven days using osmotic pumps. Aldosterone robustly increased 0.5 M NaCl intake in *Nr3c2*^lox/lox^ control mice, whereas aldosterone failed to elicit saline intake in VGLUT2-IRES-Cre::*Nr3c2*^lox/lox^ mice (**Figure 1F**). These data demonstrate that *Nr3c2* expression in VGLUT2^+^ neurons is necessary for aldosterone-induced sodium appetite and reveal a phenotype resembling that reported after HSD2 neuron ablation^3^. Next, we tested sodium deprivation and sodium depletion paradigms, which increase aldosterone secretion (**Figure S1G**) but also engage other signals associated with hypovolemia, including angiotensin II^36^. Following seven days on a sodium-deficient diet, VGLUT2-IRES-Cre::*Nr3c2*^lox/lox^ mice consumed significantly less 0.5 M NaCl than control mice (**Figure 1G**). Mutant mice also exhibited attenuated 0.5 M NaCl intake after sodium depletion by furosemide diuresis and 24 hours on a sodium-deficient diet, although to a lesser degree (**Figure 1H**). Water intake did not differ between groups in any experiment, indicating that the behavioral effects of *Nr3c2* deletion were specific to sodium appetite (**Figures 1F-1H**). Together, these data demonstrate the integral role of MR signaling in VGLUT2+ neurons for sodium appetite, and the loss of HSD2 immunoreactivity suggests that MR is required for HSD2 neuron identity, development, or survival.

### Disruption of MR in HSD2 neurons reduces their activation and sodium appetite

Since HSD2 expression was abolished in the NTS of VGLUT2-IRES-Cre::*Nr3c2*^lox/lox^ mice, this model prevented direct investigation of HSD2 neuron function following MR loss. We reasoned that CRISPR/Cas9-mediated disruption of *Nr3c2* in HSD2 neurons of adult mice would avoid the effects of developmental deletion and preserve HSD2 expression. Therefore, we developed an AAV expressing Cre-dependent Cas9 and a single guide RNA (sgRNA)^37^ targeting *Nr3c2* and used a previously developed vector targeting the non-coding Rosa26 genomic locus as a control (AAV-DIO-SaCas9-U6-sg*Nr3c2* and AAV-DIO-SaCas9-U6-sg*Rosa26*, respectively). These vectors were injected into the NTS of *Hsd11b2*-IRES-Cre mice to achieve HSD2 neuron-specific disruption of *Nr3c2* (HSD2^Cas9-sg*Nr3c2*^ or HSD2^Cas9-sg*Rosa*26^; **Figure 2A**). To assess the efficacy of our Cas9 vectors, we collected HSD2 neuron nuclei expressing either Cas9 AAV and sequenced the genomic regions targeted by sg*Rosa26* or sg*Nr3c2* (**Figure S2A**). Approximately 50% of sg*Rosa26*-targeted reads contained mutations (25% deletions, 17.6% insertions, and 7% base pair substitutions), consistent with previous findings^37^. In comparison, ∼75% of sg*Nr3c2*-targeted reads contained mutations (71.5% deletions, 1.3% insertions, and 4.3% base pair substitutions), confirming efficient editing of the *Nr3c2* locus (**Figure S2B**). Importantly, unlike developmental MR deletion, adult disruption of *Nr3c2* preserved HSD2 immunoreactivity (**Figure 2B**), allowing us to investigate the functional consequences of MR loss in mature HSD2 neurons.

**Figure 2:**
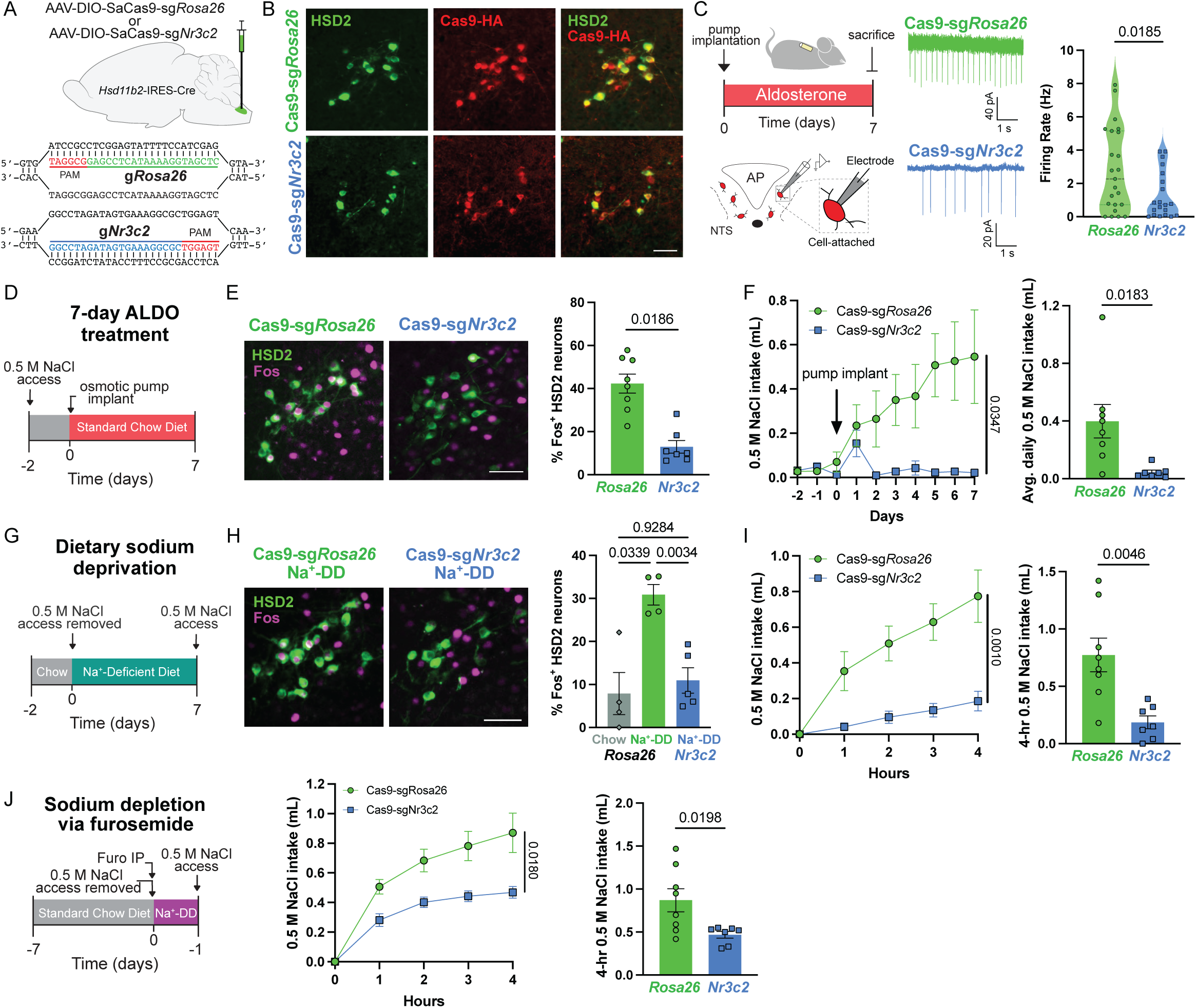
CRISPR-Cas9 disruption of *Nr3c2* in HSD2 neurons severely impairs sodium appetite. **A)** Schematic showing injection of Cre-dependent AAV-Cas9 targeting either *Rosa26* or *Nr3c2* into the NTS of *Hsd11b2*-IRES-Cre mice (top) and sgRNA+PAM sequences (bottom). **B)** Representative brain sections showing HSD2 immunofluorescence (IF; left), Cas9-HA IF (middle), and both (right) in HSD2^Cas9-sg*Rosa*26^ (top) or HSD2^Cas9-sg*Nr3c2*^ (bottom) mice (scale bar = 50 μm). **C)** Schematic for electrophysiology recordings after aldosterone treatment in *Hsd11b2*-IRES-Cre mice injected with Cre-dependent AAV-Cas9 vectors (left), with representative cell-attached recordings of HSD2 neuron action potential firing (middle), and mean firing rate (right) from HSD2^Cas9-sg*Rosa*26^ or HSD2^Cas9-sg*Nr3c2*^ mice following 7 days of aldosterone treatment (n = 20-23 neurons from 2-3 mice/group). Data are represented as median and quartiles. **D)** Schematic describing the aldosterone-induced sodium appetite model. **E)** Representative images showing c-Fos expression in HSD2 neurons of HSD2^Cas9-sg*Rosa*26^ (left) or HSD2^Cas9-sg*Nr3c2*^ (middle) mice following 7 days of aldosterone treatment and the percentage of HSD2 neurons expressing c-Fos (right; scale bar = 50 μm). **F)** 0.5 M NaCl intake in HSD2^Cas9-sg*Rosa*26^ or HSD2^Cas9-sg*Nr3c2*^ mice during 7 days of aldosterone treatment (left) and average 0.5 M NaCl intake per day (right; n = 7-8 mice/group). Data are represented as mean ± SEM. ALDO = aldosterone. **G)** Schematic describing the dietary sodium deprivation sodium appetite model. **H)** Representative images showing c-Fos expression in HSD2 neurons of HSD2^Cas9-sg*Rosa*26^ (left) or HSD2^Cas9-sg*Nr3c2*^ (middle) mice following dietary sodium deprivation and the percentage of HSD2 neurons expressing c-Fos (right; n = 4-5 mice/group; scale bar = 50 μm). **I)** 0.5 M NaCl intake in HSD2^Cas9-sg*Rosa*26^ or HSD2^Cas9-sg*Nr3c2*^ mice over 4 hours following dietary sodium deprivation (left) and total 0.5 M NaCl intake in 4 hours (right; n = 7-8 mice/group). Data are represented as mean ± SEM. **J)** Schematic describing the furosemide-induced sodium depletion sodium appetite model (left), 0.5 M NaCl intake in HSD2^Cas9-sg*Rosa*26^ or HSD2^Cas9-sg*Nr3c2*^ mice over 4 hours following sodium depletion via furosemide (middle) and total 0.5 M NaCl intake in 4 hours (right; n = 7-8 mice/group). Furo = furosemide; Na^+^-DD = Na^+^-deficient diet. Statistical analysis of saline intake over time between groups was performed using a two-way repeated-measures ANOVA with Geisser-Greenhouse correction. Two-group comparisons without repeated-measures were performed with a Welch’s t-test, and three-group comparisons were performed using one-way Brown-Forsythe and Welch ANOVA tests followed by Dunnett T3 correction. p < 0.05 was considered statistically significant.

Given that aldosterone activates HSD2 neurons, we first asked whether *Nr3c2* disruption impairs this response. Indeed, following aldosterone treatment, HSD2^Cas9-sg*Nr3c2*^ mice exhibited significant reductions in firing rate and c-Fos expression in HSD2 neurons compared to HSD2^Cas9-sg*Rosa*26^ controls (**Figures 2C-2E**). We next assessed whether disruption of *Nr3c2* in HSD2 neurons alters sodium appetite. As expected, aldosterone treatment robustly increased 0.5 M NaCl intake in HSD2^Cas9-sg*Rosa*26^ mice, but this response was abolished in HSD2^Cas9-sg*Nr3c2*^ mice (**Figure 2F**) with no difference in water intake between groups (**Figures S2C and S2D**). Of interest, HSD2^Cas9-sg*Nr3c2*^ mice also exhibited impaired sodium deprivation-induced c-Fos expression in HSD2 neurons (**Figures 2G and 2H**), indicating that MR signaling contributes to HSD2 neuron activation during sodium deficiency. Consistent with this impaired activation, HSD2^Cas9-sg*Nr3c2*^ mice drank significantly less 0.5 M NaCl than HSD2^Cas9-sg*Rosa*26^ controls following both dietary sodium deprivation and sodium depletion via furosemide diuresis (**Figures 2I and 2J**) with no difference in water intakes between groups (**Figures S2E and S2F**). No differences in 0.5 M NaCl intake were observed between HSD2^Cas9-sg*Rosa*26^, HSD2^mCherry^, or Cre-negative Cas9-sg*Nr3c2*-injected controls in any of the three paradigms (**Figures S2G-S2J**). Notably, HSD2^Cas9-sg*Nr3c2*^ mice exhibited a sodium appetite phenotype remarkably similar to that of VGLUT2-IRES-Cre::*Nr3c2*^lox/lox^ mice, indicating that loss of MR specifically in HSD2 neurons largely accounts for the behavioral phenotype observed following its broader deletion from VGLUT2^+^ neurons. Together, these data demonstrate that aldosterone-MR signaling in HSD2 neurons is a critical driver of their activity and important for sodium appetite during sodium deficiency and mineralocorticoid excess.

### Aldosterone selectively drives a synaptic transcriptional program in HSD2 neurons

HSD2 neurons are the only neurons in the adult mouse brain that coexpress HSD2 and MR^12,13^, providing a unique population in which to investigate the effects of aldosterone-MR signaling on neuronal function. Since MR is a steroid hormone receptor that regulates gene expression, we hypothesized that aldosterone drives a transcriptional program that enhances HSD2 neuron activity to promote sodium appetite. However, the limited number of HSD2 neurons (∼200 neurons per mouse^3,12,14^) makes direct mapping of MR-DNA interactions challenging. In addition, gene expression changes identified in our previous scRNA-seq study of HSD2 neurons after eight days of sodium deprivation cannot be specifically attributed to aldosterone, as sodium deficiency engages additional signals that activate HSD2 neurons, and sustained neuronal activity itself can alter gene expression^38^. We therefore performed HSD2 neuron-enriched snRNA-seq of the caudal NTS to decipher the transcriptional response to aldosterone and identify candidate molecular mechanisms regulating HSD2 neuron function.

*Hsd11b2*-IRES-Cre mice received NTS injections of AAV-DIO-H2B-mCherry to drive nuclear mCherry expression in HSD2 neurons. Mice were then implanted with osmotic pumps delivering vehicle or aldosterone for 24 hours. This treatment produced robust c-Fos expression in HSD2 neurons, comparable to seven days of aldosterone administration, and increased their action potential firing (**Figures 3A and 3B**)^3,14^. After tissue collection, we used fluorescence-activated nuclei sorting (FANS) to enrich for mCherry^+^ HSD2 neuron nuclei (**Figure 3C**), followed by sequencing and analysis using Seurat^39^ in R. The resulting dataset contained 22,217 nuclei that passed quality control filtering, with cell types throughout the dorsomedial medulla represented (**Figures 3D and S3A; Table S1A**), and an average read depth of 23,000 reads per neuronal nucleus. Next, we subclustered all neuronal nuclei (*Rbfox3*^+^ and *Tubb3*^+^; **Figure S3B**) and subsequently analyzed glutamatergic, GABAergic, and cholinergic populations separately, yielding 5,317 glutamatergic (*Slc17a6^+^*), 3,384 GABAergic (*Slc32a1^+^*), and 437 cholinergic (*Chat^+^*) neuronal nuclei (**Figures 3E**, **3F, and S3C-S3I**). Marker genes were identified for each cluster within these three broad neuronal subtypes, many of which have been described previously (**Tables S1B-S1D**)^40–42^. Importantly, the largest glutamatergic cluster (GLUT1.Hsd11b2) exhibited strong and specific *Hsd11b2* expression, confirming successful enrichment for HSD2 neurons (**Figure 3G**).

**Figure 3:**
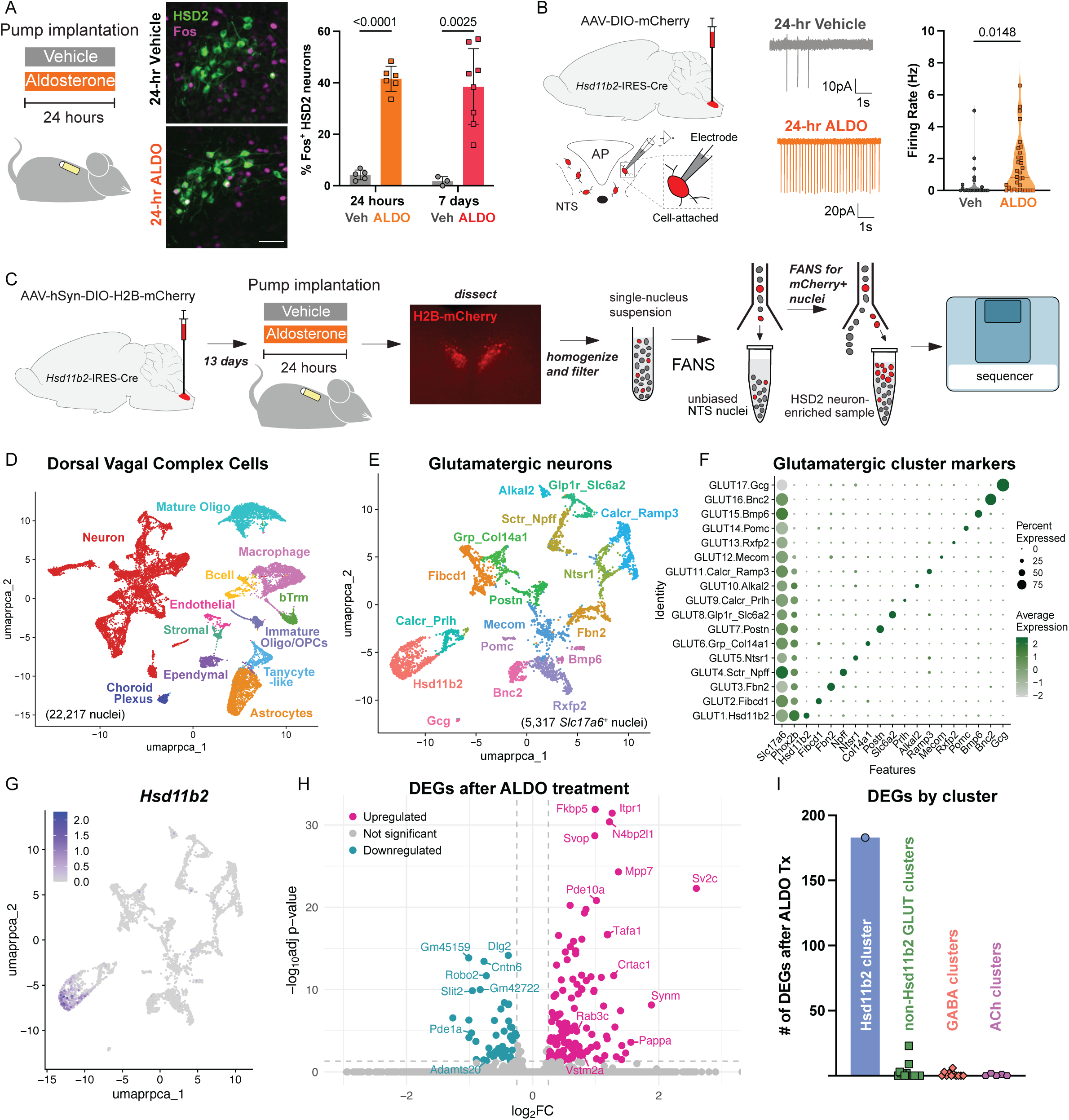
HSD2 neuron-enriched snRNA-seq of the caudal NTS following aldosterone treatment. **A)** Schematic describing 24-hour aldosterone treatment experiments (left), representative brain sections showing HSD2 and c-Fos immunofluorescence (IF) following 24-hour vehicle (middle-top) or aldosterone (middle-bottom) treatment, and the percentage of HSD2 neurons expressing c-Fos following 24 hours or 7 days of vehicle or aldosterone treatment (right; 24-hr treatment, n = 5-6 mice/group; 7-day treatment, n = 3 mice vehicle and n = 8 mice aldosterone). Data are represented as mean ± SEM and were analyzed with unpaired t-tests corrected for multiple comparisons (scale bar = 50 μm). **B)** Schematic for electrophysiology recordings after aldosterone treatment in *Hsd11b2*-IRES-Cre mice injected with Cre-dependent AAV-mCherry (left), with representative cell-attached recordings of HSD2 neuron action potential firing (middle), and mean firing rate (right) from 24-hour treatment with vehicle or aldosterone (ALDO; n = 22-32 neurons from 2 mice/group). Data are represented as median and quartiles and were analyzed using a Welch’s t-test. **C)** Schematic of snRNA-seq experiment workflow describing aldosterone treatment and enrichment for HSD2 neurons within the NTS sample. **D)** UMAP of all dorsal vagal complex cells, comprised of 22,217 nuclei. **E)** UMAP of all glutamatergic neurons (*Slc17a6*^+^), comprised of 5,317 nuclei. **F)** Dot plot showing *Slc17a6*^+^ cluster markers. **G)** Feature plot depicting the expression of *Hsd11b2* in *Slc17a6*^+^ neurons. **H)** Volcano plot showing DEGs in the GLUT1.Hsd11b2 cluster. Downregulated genes shown in blue and upregulated genes shown in pink. DEGs, differentially expressed genes. Absolute log_2_ fold change (log_2_FC) > 0.25 and adjusted p < 0.05 were considered statistically significant. **I)** Number of DEGs after aldosterone treatment for each cluster in the caudal NTS snRNA-seq experiment, grouped by neurotransmitter type.

Analysis of the GLUT1.Hsd11b2 cluster identified several enriched marker genes, including *Slc9a3*, *Tspan11*, and *Ptger2* (**Figure S4A; Table S1B**). *Slc9a3* encodes the Na+/H+ exchanger NHE3, which is vital for renal sodium reabsorption^43,44^, and we confirmed its expression in HSD2 neurons using fluorescence *in situ* hybridization (FISH; **Figure S4B**). Notably, aldosterone treatment altered the expression of 183 genes in the GLUT1.Hsd11b2 cluster (**Figures 3H and S4C; Table S2**). In contrast, most other neuronal clusters exhibited no DEGs, and only one had more than 10 (GLUT15.Bmp6; 23 DEGs; **Figure 3I; Tables S2A-S2C**). Of the 183 DEGs in HSD2 neurons, only *Nrg3os* was also differentially expressed in another glutamatergic cluster (GLUT2.Fibcd1). Thus, aldosterone elicits a remarkably selective transcriptional response in HSD2 neurons compared with neighboring neuronal populations.

We next examined whether aldosterone-regulated genes in the GLUT1.Hsd11b2 cluster included established MR targets. Many canonical MR-responsive genes were either not detected (e.g., *Tsc22d3*, formerly “*Gilz*”) or unchanged following aldosterone treatment (e.g., *Sgk1*)^28,45^. HSD2 neurons also lacked expression of epithelial sodium channel (ENaC) subunits, in agreement with prior work^14^. Nevertheless, aldosterone increased the expression of some established MR-regulated genes, including *Fkbp5*, *Zbtb16*, and *Atp1a1* (**Figure S4D**)^28,45–47^. Of interest, *Fkbp5* encodes FK506 binding protein 5, a steroid receptor cochaperone that participates in negative feedback regulation of MR and glucocorticoid receptor (GR) signaling^48,49^. Consistent with its regulation by MR, *Fkbp5* expression decreases following MR knockdown in the hippocampus^50,51^. The function of the other MR-regulated DEGs in neurons is less well understood. Together, these data are consistent with cell type-specific MR-associated transcriptional programs^27,52^ and indicate that neuronal responses to aldosterone differ substantially from canonical MR signaling in epithelial tissues.

Because sodium deficiency increases aldosterone secretion, we expected overlap between the transcriptional responses to aldosterone treatment and sodium deprivation^14^. Surprisingly, only 20 DEGs were shared between the two datasets (**Figures S4E and S4F**). Notable shared DEGs included upregulation of *Cacna1i*, encoding the T-type calcium channel Ca_v_3.3; *Kcnip1,* encoding voltage-gated potassium channel interacting protein 1; and the MR-regulated gene *Atp1a1* referred to above. In contrast, the postsynaptic genes *Nlgn1* and *Dlg2*^53,54^ showed opposing regulation, with both downregulated by aldosterone but upregulated by sodium deprivation. This discrepancy may reflect distinct adaptations in the postsynaptic organization of HSD2 neurons associated with mineralocorticoid excess and sodium deficiency, respectively. Thus, the transcriptional response to 24 hours of aldosterone treatment differs substantially from that induced by prolonged sodium deprivation, potentially reflecting differences in treatment duration, additional physiological signals, and persistent neuronal activation during sodium deficiency.

To identify biological systems associated with aldosterone-responsive genes in HSD2 neurons, we performed gene ontology (GO) enrichment analysis of the DEGs. Numerous enriched terms were associated with synaptic function, including synapse organization, neuron projection organization, and synaptic vesicles (**Figure S4G**). Aldosterone altered the expression of genes associated with both pre- and postsynaptic function, including upregulation of *Grik4* and *Gria1*, which encode kainate and AMPA receptor subunits, respectively, and could enhance excitatory synaptic responsiveness in HSD2 neurons. Genes involved in neuronal and synaptic organization were also differentially expressed, including upregulation of lateral olfactory tract usher substance (LOTUS/*Crtac1*)^55^ and the microtubule-associated protein 1B (*Map1b*)^56^. Conversely, Roundabout 2 (*Robo2*) and SLIT 2 (*Slit2*), which encode binding partners that negatively regulate synapse assembly^57^, were downregulated. Collectively, these findings identify an aldosterone-induced transcriptional program enriched for genes associated with synaptic organization and function, suggesting that aldosterone-MR signaling may enhance synaptic transmission from HSD2 neurons.

### Aldosterone enhances synaptic transmission during repeated activation of HSD2 neurons

The preponderance of synaptic DEGs prompted us to investigate whether aldosterone enhances synaptic transmission from HSD2 neurons to downstream targets. We focused on HSD2 neuron projections to the ventrolateral bed nucleus of the stria terminalis (BSTvL), as stimulation of these projections has been shown to drive sodium intake^14^. ChrimsonR-tdTomato^58^ was expressed in HSD2 neurons, and animals were subsequently treated with vehicle or aldosterone prior to whole-cell patch-clamp recordings of BSTvL neurons from *ex vivo* brain slices (**Figure 4A**). To assess changes in HSD2 neuron synaptic function, we recorded light-evoked excitatory postsynaptic currents (le-EPSCs) using paired-pulse stimulation in the presence of tetrodotoxin (TTX) and 4-aminopyridine (4-AP) to isolate monosynaptic connections (**Figure 4B**). HSD2 à BSTvL recordings exhibited approximately 70% connectivity regardless of treatment (**Figure 4C**). Importantly, aldosterone did not alter the amplitude of the first le-EPSC but increased the second response, resulting in an increased paired-pulse ratio (PPR; **Figures 4D-4F**), without significant changes in response latency, decay time, or charge (**Figures S5A-S5C**). These data demonstrate that aldosterone reduces paired-pulse depression at HSD2 neuron efferent synapses, consistent with an enhanced capacity to maintain neurotransmitter release during repeated activation.

**Figure 4:**
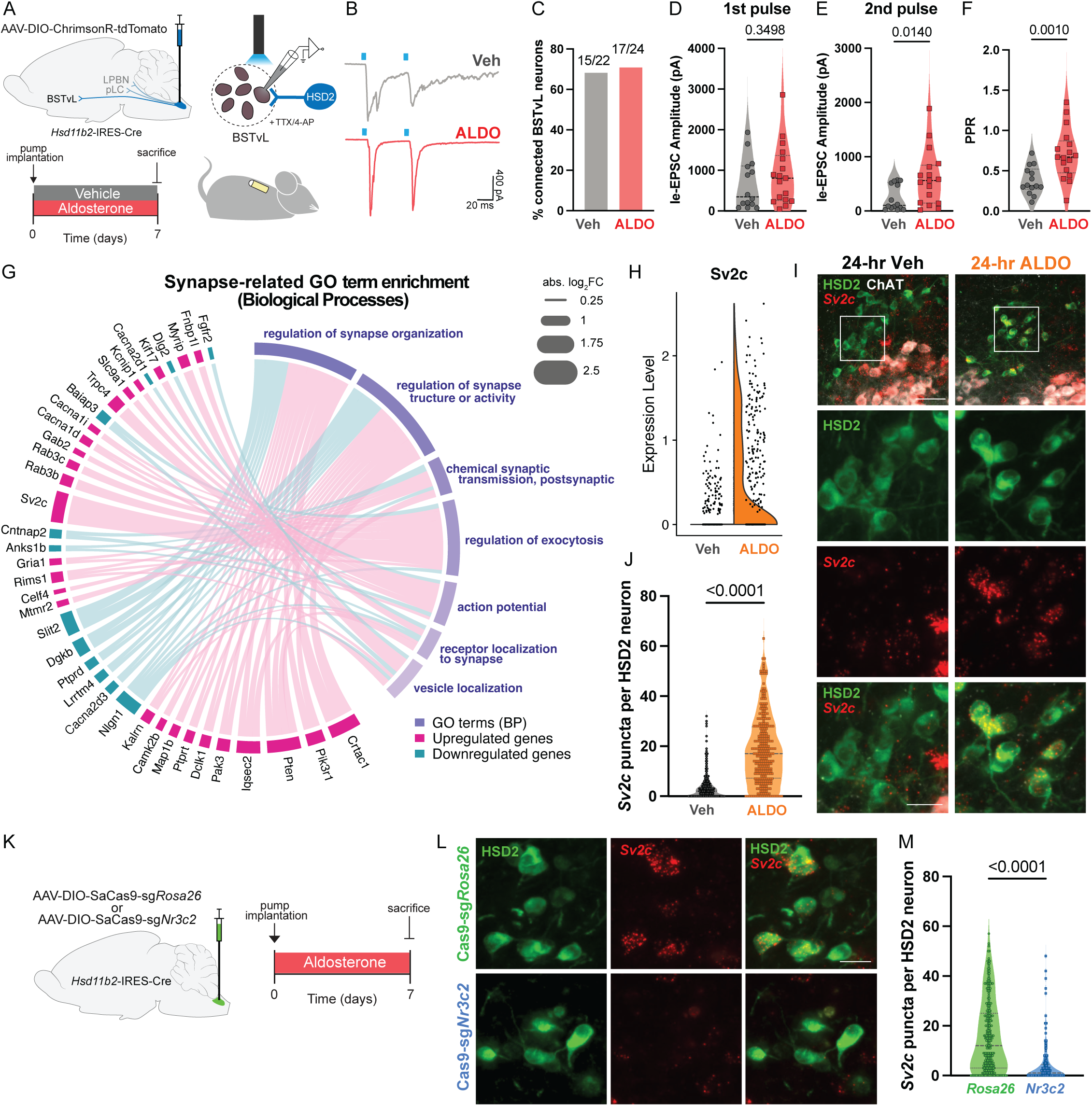
Aldosterone enhances neurotransmission during repeated activation and drives a synapse-associated transcriptional program. **A)** Schematic showing injection of Cre-dependent AAV-ChrimsonR-tdTomato into the NTS of *Hsd11b2*-IRES-Cre mice and workflow for electrophysiology recordings in the BSTvL after 7-day vehicle or aldosterone treatment. **B)** Representative light-evoked paired-pulse EPSCs in BSTvL neurons from mice treated with vehicle (top) or aldosterone (ALDO; bottom). **C)** Connection rate of HSD2 neurons à BSTvL (n = 22-24 neurons from 2-4 mice/group). **D)** Light-evoked EPSC response amplitude to the first pulse (n = 14-17 neurons from 2-4 mice/group). Data are represented as median and quartiles. **E)** Light-evoked EPSC response amplitude to the second pulse (n = 14-17 neurons from 2-4 mice/group). Data are represented as median and quartiles. **F)** Paired-pulse ratio (PPR) of light evoked-EPSC response amplitudes (n = 14-17 neurons from 2-4 mice/group). Data are represented as median and quartiles. **G)** Circos plot depicting top 7 synapse-related GO terms simplified by semantic similarity and the differentially expressed genes (DEGs) contributing to those terms. **H)** Split violin plot showing expression of *Sv2c* in cluster GLUT1.Hsd11b2 from snRNA-seq of vehicle- and aldosterone-treated mice. **I)** Representative brain sections from mice treated with vehicle (left) or aldosterone (right) for 24 hours showing in descending order: low-magnification view of dorsal vagal complex *Sv2c* fluorescence *in situ* hybridization (FISH) colocalized with HSD2 and ChAT immunofluorescence (IF) (scale bar = 50 μm); - magnification view of HSD2 IF in boxed regions, high-magnification view of *Sv2c* FISH in boxed regions, and high-magnification view of both HSD2 IF and *Sv2c* FISH (scale bar = 20 μm). **J)** Total number of *Sv2c* puncta per HSD2 neuron (n = 284-296 neurons from 3 mice/group). **K)** Schematic showing injection of Cre-dependent AAV-Cas9 targeting either *Rosa26* or *Nr3c2* into the NTS of *Hsd11b2*-IRES-Cre mice (left) and schematic describing the aldosterone treatment model (right). **L)** Representative brain sections from HSD2^Cas9-sg*Rosa*26^ (top) and HSD2^Cas9-sg*Nr3c2*^ (bottom) mice following 7 days of aldosterone treatment showing, HSD2 immunofluorescence (left; IF), *Sv2c* fluorescence *in situ* hybridization (middle; FISH), and both (right; scale bar = 20 μm). **M)** Total number of *Sv2c* puncta per HSD2 neuron (n = 153-245 neurons from 3 mice/group). Data are represented as median and quartiles. All statistical analyses were performed using a Welch’s t-test. p < 0.05 was considered statistically significant.

To identify molecular mechanisms that could underlie aldosterone-induced changes in synaptic transmission, we returned to the snRNA-seq data and examined DEGs contributing to GO Biological Processes terms related to “synapses”, “exocytosis”, “vesicles”, and “action potential” (**Figure 4G**). Expression of *Cacna2d1* and *Cacna2d3*, which encode voltage-gated calcium channel α2δ-1 and α2δ-3 auxiliary subunits, respectively, were decreased. α2δ subunits regulate channel trafficking and function^59^, suggesting that their downregulation could alter presynaptic calcium influx and vesicle release dynamics, potentially contributing to reduced paired-pulse depression. Concurrently, several genes associated with the presynaptic active zone and synaptic vesicles were upregulated, including regulating synaptic membrane exocytosis protein 1 (*Rims1*)^60^, Ras-related protein Rab-3C (*Rab3c*)^61^, and synaptic vesicle glycoprotein 2 C (*Sv2c*)^62^. These genes encode components of presynaptic machinery involved in vesicle docking, priming, and release, potentially contributing to the observed reduction in paired-pulse depression. Notably, *Sv2c* exhibited the largest aldosterone-induced increase among all DEGs in the GLUT1.Hsd11b2 cluster (∼6-fold; **Figure 4H**). Independent FISH quantification in HSD2 neurons confirmed an ∼5-fold increase in *Sv2c* expression after 24 hours of aldosterone treatment (**Figures 4I and 4J**). Moreover, disruption of *Nr3c2* in HSD2 neurons prevented the aldosterone-induced increase in *Sv2c* expression, demonstrating that this response requires MR (**Figures 4K-4M**).

### Disruption of *Sv2c* in HSD2 neurons impairs sodium appetite

SV2C belongs to the synaptic vesicle glycoprotein 2 family and, together with SV2A and SV2B, SV2 proteins are integral synaptic vesicle membrane proteins implicated in vesicle loading, priming and docking, and calcium-dependent neurotransmitter release^63,64^. Given its role in synaptic vesicle function and robust induction in HSD2 neurons by aldosterone-MR signaling, we hypothesized that SV2C is important for aldosterone-mediated salt appetite. To test this, we employed the same Cre-dependent CRISPR-Cas9 strategy to disrupt *Sv2c* specifically in HSD2 neurons (HSD2^Cas9-sg*Sv2c*^ or HSD2^Cas9-sg*Rosa*26^; **Figures 5A and 5B**). HSD2^Cas9-sg*Sv2c*^ mice drank significantly less 0.5 M NaCl than HSD2^Cas9-sg*Rosa*26^ mice during the course of aldosterone treatment (**Figures 5C and 5D**). Despite reduced sodium intake, aldosterone-induced c-Fos expression did not differ between HSD2^Cas9-sg*Rosa*26^ and HSD2^Cas9-sg*Sv2c*^ mice (**Figure 5E**), suggesting that *Sv2c* disruption does not prevent aldosterone-mediated activation of HSD2 neurons. No significant differences in saline intake were observed following dietary sodium deprivation or furosemide-induced sodium depletion in HSD2^Cas9-sg*Sv2c*^ mice (**Figures 5F and 5G**), and water intake did not differ between groups in any experiments (**Figures S6A-S6D**).

**Figure 5:**
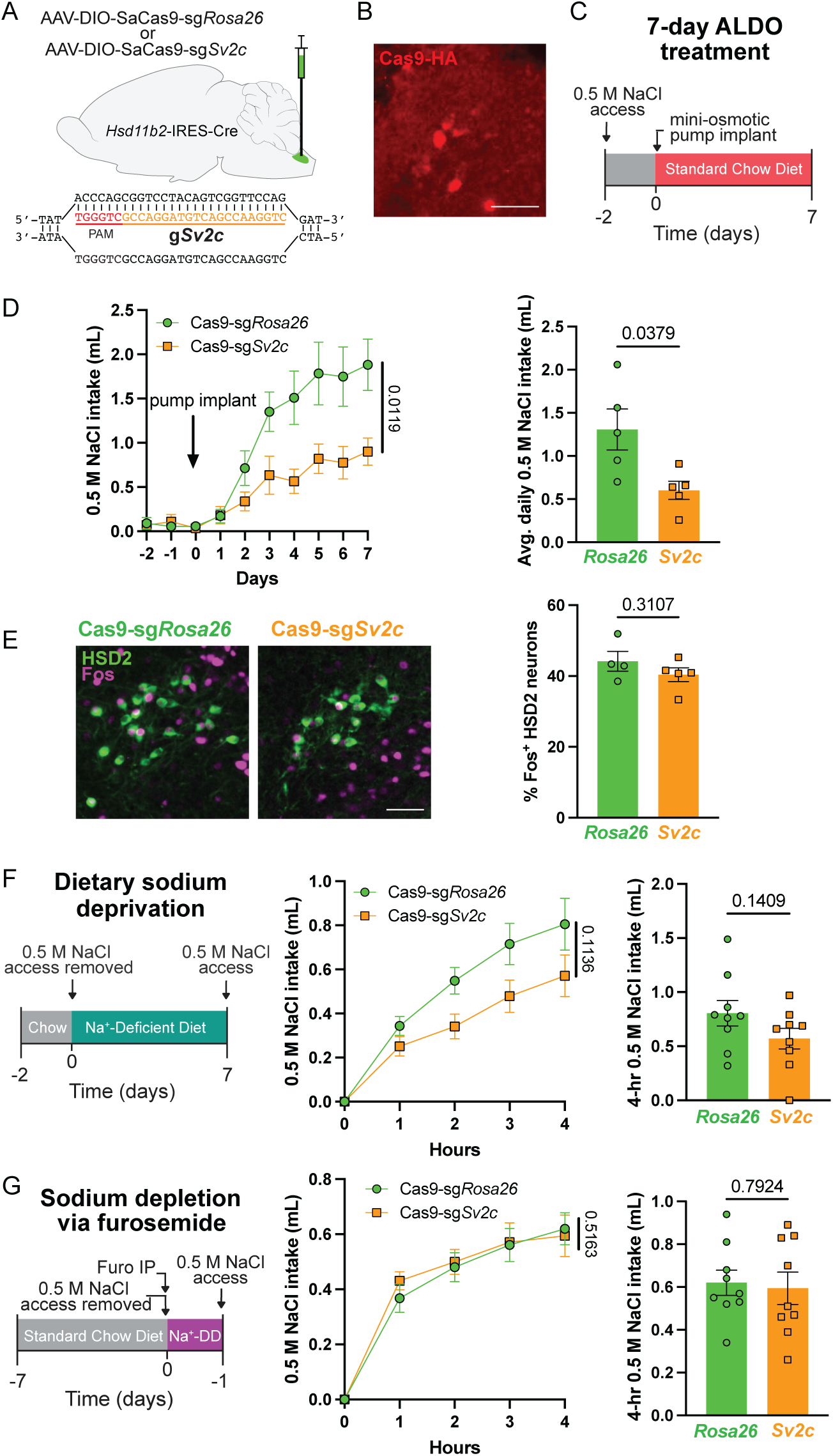
Disruption of *Sv2c* in HSD2 neurons blunts aldosterone-induced saline intake. **A)** Schematic showing injection of Cre-dependent AAV-Cas9 targeting either *Rosa26* or *Sv2c* into the NTS of *Hsd11b2*-IRES-Cre mice (top) and sgRNA+PAM sequence for *Sv2c* (bottom). **B)** Representative brain section showing Cas9-HA immunofluorescence (IF) in HSD2^Cas9-sg*Sv2c*^ mice (scale bar = 50 μm). **C)** Schematic describing the aldosterone-induced sodium appetite model. ALDO = aldosterone. **D)** 0.5 M NaCl intake in HSD2^Cas9-sg*Rosa*26^ or HSD2^Cas9-sg*Sv2c*^ mice during 7 days of aldosterone treatment (left) and average 0.5 M NaCl intake per day (right; n = 5 mice/group). **E)** Representative images showing c-Fos expression in HSD2 neurons of HSD2^Cas9-sg*Rosa*26^ (left) or HSD2^Cas9-sg*Sv2c*^ (middle) mice following 7 days of aldosterone treatment and the percentage of HSD2 neurons expressing c-Fos (right; n = 4-5 mice/group; scale bar = 50 μm). **F)** Schematic describing the dietary sodium deprivation sodium appetite model (left), with 0.5 M NaCl intake in HSD2^Cas9-sg*Rosa*26^ or HSD2^Cas9-sg*Sv2c*^ mice over 4 hours following dietary sodium deprivation (middle) and total 0.5 M NaCl intake over 4 hours (right; n = 9 mice/group). **G)** Schematic describing the furosemide-induced sodium depletion sodium appetite model (left), 0.5 M NaCl intake in in HSD2^Cas9-sg*Rosa*26^ or HSD2^Cas9-sg*Sv2c*^ mice over 4 hours following sodium depletion via furosemide (middle), total 0.5 M NaCl intake in 4 hours (right; n = 9 mice/group). Furo = furosemide; Na^+^-DD = Na^+^-deficient diet. All data are represented as mean ± SEM. Statistical analysis of saline intake over time between groups was performed using a two-way repeated-measures ANOVA with Geisser-Greenhouse correction. All other statistical analyses were performed with a Welch’s t-test. p < 0.05 was considered statistically significant.

An independent approach using short hairpin RNA (shRNA)-mediated knockdown of *Sv2c* in HSD2 neurons also reduced sodium appetite, although this manipulation produced a more pronounced behavioral phenotype, including attenuated deprivation-induced saline intake and aldosterone-induced c-Fos expression (**Figures S6E-S6K**). Finally, Cre-dependent overexpression of SV2C in HSD2 neurons (AAV-DIO-*Sv2c*-GFP) did not increase saline intake compared to GFP controls (AAV-DIO-eGFP; **Figures S6L-S6P**), indicating that increased *Sv2c* expression alone is insufficient to drive sodium appetite. Collectively, these findings identify *Sv2c* as an aldosterone-MR-regulated mediator of sodium appetite driven by HSD2 neurons.

## Discussion

Primary aldosteronism is the most common endocrine cause of secondary hypertension and accounts for approximately 20% of treatment-resistant hypertension^65,66^. High salt intake in these patients exacerbates deleterious cardiovascular remodeling and kidney damage^67,68^. Yet, patients with primary aldosteronism exhibit high dietary sodium intake^69,70^, which may be driven, in part, by the effects of aldosterone on salt appetite. In this study, we investigated the neuromolecular mechanisms through which aldosterone drives sodium appetite. Loss of MR in HSD2 neurons impaired their activation and sodium appetite, completely abolishing aldosterone-induced saline intake. Profiling of the caudal NTS following aldosterone treatment revealed a remarkably selective transcriptional response in HSD2 neurons, with DEGs enriched for synaptic function. Consistent with this, optogenetic paired-pulse recordings demonstrated that aldosterone reduces paired-pulse depression at HSD2 neuron synapses, suggesting an enhanced capacity to maintain synaptic transmission during repeated stimulation. Among the genes upregulated by aldosterone, *Sv2c* exhibited the greatest increase in expression, and its selective disruption in HSD2 neurons diminished aldosterone-induced saline intake. These results establish that aldosterone-MR signaling in HSD2 neurons is essential for aldosterone-induced sodium appetite and implicate transcriptional regulation of synaptic machinery as a mechanism through which aldosterone augments HSD2 neuron function to promote sodium ingestion.

### MR regulation of HSD2 neuron identity and function

Both MR-dependent and MR-independent mechanisms across multiple brain regions have been proposed to mediate the effects of aldosterone on sodium appetite^71–74^. However, because circulating glucocorticoid concentrations greatly exceed those of aldosterone, selective activation of MR by aldosterone requires HSD2-mediated glucocorticoid inactivation^9–12^. HSD2 neurons in the NTS are the only neurons in the adult mouse brain known to express both HSD2 and MR^12,13^, uniquely positioning them to directly sense circulating aldosterone. Consistent with this, ablation of HSD2 neurons abolishes aldosterone-induced sodium appetite^3^. The present work extends these findings by demonstrating that disruption of *Nr3c2* specifically in adult HSD2 neurons abolishes aldosterone-induced saline intake and impairs deficiency-induced sodium ingestion. Notably, both saline intake and HSD2 neuron c-Fos immunoreactivity were significantly reduced in HSD2^Cas9-sg*Nr3c2*^ mice following sodium deprivation, suggesting that aldosterone-MR signaling contributes substantially to HSD2 neuron activity during sodium deficiency. Thus, MR signaling within HSD2 neurons is essential for aldosterone-induced sodium appetite and contributes to sodium intake during sodium deficiency.

Remarkably, developmental deletion of *Nr3c2* from VGLUT2^+^ neurons completely abolished detectable HSD2 immunoreactivity in the NTS and strongly reduced sodium appetite. Although unexpected, previous studies have reported apparent increases in HSD2 expression following mineralocorticoid treatment in multiple cell types^75,76^, including in HSD2 neurons^6^, and MR deletion markedly reduces *Hsd11b2* expression in the epidermis^77^, suggesting that MR signaling can regulate HSD2 expression. In contrast, HSD2 expression persisted following CRISPR-Cas9-mediated disruption of *Nr3c2* in adult HSD2 neurons. Together, these findings raise the possibility that MR signaling may be required during development for establishment of HSD2 expression, HSD2 neuron development, or survival, but disruption of MR in adulthood does not eliminate HSD2 expression. The alternative molecular markers for HSD2 neurons identified in this study may enable future studies to distinguish whether developmental MR deletion causes selective loss of *Hsd11b2* expression or instead disrupts HSD2 neuron development or survival.

### Neuromolecular mechanisms of aldosterone-MR signaling

Consistent with the requirement of HSD2 for conferring aldosterone sensitivity, snRNA-seq of the caudal NTS following aldosterone treatment revealed selective differential gene expression within the GLUT1.Hsd11b2 cluster. Although snRNA-seq cannot distinguish direct MR transcriptional targets from secondary changes in gene expression, these data provide insight into the transcriptional response driven by aldosterone-MR signaling in neurons. Multiple established MR-responsive genes were upregulated, including *Fkbp5*^28,45,50,51^, *Zbtb16*^28,78^, and *Atp1a1*. Among these, *Zbtb16* encodes an aldosterone-responsive transcription factor implicated in renal ENaC regulation^79^, which is also involved in glucocorticoid receptor-mediated cortical neurogenesis^80^ and control of energy homeostasis by hypothalamic neurons^81^. Despite regulation of these established MR targets, the broader transcriptional response to aldosterone in HSD2 neurons differed substantially from that described in the kidney and other aldosterone-sensitive tissues. Instead, aldosterone-responsive genes were enriched for pathways associated with synaptic function and remodeling. Some of these changes may reflect increased neuronal activity rather than direct MR-mediated transcription. Nevertheless, our snRNA-seq data reveal an unexpected synapse-associated transcriptional signature in HSD2 neurons that was not prominent following sodium deprivation, raising the possibility that similar programs may be engaged by MR signaling in other neuronal populations.

Prior scRNA-seq of HSD2 neurons following prolonged sodium deprivation identified substantial differential expression of voltage-gated ion channels, including hyperpolarization-activated cyclic nucleotide-gated (HCN) channels, the “cardiac” sodium channel *Scn5a*/Na_v_1.5, and the R-type calcium channel *Cacna1e*/Ca_v_2.3. These channels were subsequently shown to cooperatively regulate pacemaker-like intrinsic activity of HSD2 neurons during sodium deficiency. Similar ion channel changes were not prominent following 24 hours of aldosterone treatment, suggesting that the increased intrinsic activity under these conditions may arise through distinct molecular mechanisms. Despite robust HSD2 neuron activation under both conditions, only 20 DEGs were shared between the aldosterone- and sodium deprivation-induced transcriptional responses. Direct comparison of these datasets is limited, however, by differences in treatment duration and sequencing methodology: the present study used snRNA-seq, while the sodium deprivation study employed Smart-seq2 of a much smaller number of manually isolated HSD2 neurons. Nevertheless, the distinct transcriptional signatures are consistent with sodium deprivation integrating additional signals associated with sodium deficiency and hypovolemia beyond aldosterone alone, such as angiotensin II and neural inputs.

The enrichment of aldosterone-responsive genes associated with synaptic function prompted us to investigate neurotransmission from HSD2 neurons to downstream targets. Optically evoked paired-pulse recordings from BSTvL neurons receiving HSD2 input revealed reduced paired-pulse depression after aldosterone treatment, driven by a selective increase in the response to the second light pulse. This reduction in paired-pulse depression could reflect several presynaptic mechanisms, including accelerated vesicle replenishment or prolonged presynaptic calcium signaling. Such adaptations could facilitate the maintenance of neurotransmitter release onto downstream targets during periods of elevated HSD2 neuron firing induced by aldosterone and sodium deficiency. Among the transcriptional changes in HSD2 neurons, *Sv2c* was particularly notable for exhibiting the largest aldosterone-induced increase and encoding an integral synaptic vesicle protein implicated in vesicle trafficking and exocytosis. Thus, increased *Sv2c* expression provides a candidate mechanism through which aldosterone could alter neurotransmitter release from HSD2 neurons. Importantly, disruption of *Sv2c* in HSD2 neurons reduced saline intake, demonstrating a functional role for SV2C in aldosterone-induced sodium appetite.

SV2C has the most restricted expression pattern compared to the other members of its family (SV2A and SV2B), with expression noted in the basal ganglia and hindbrain^62,63^. Despite being an integral component of the synaptic vesicle^82^, the precise function of SV2 proteins in neuronal transmission remains unclear, although they are thought to regulate multiple aspects of synaptic vesicle trafficking and calcium-dependent exocytosis. Thus, SV2C in HSD2 neurons could influence neurotransmitter release through effects on vesicle priming^83,84^, vesicle recruitment to the plasma membrane^85^, or binding calcium and synaptotagmin^83,86–88^. Moreover, other aldosterone-responsive genes, including *Rims1*, *Cacna2d1*, and *Cacna2d3*, could also independently alter presynaptic calcium dynamics or release probability^89,90^. Because residual presynaptic calcium following an initial stimulus can augment subsequent neurotransmitter release^91^, coordinated changes in vesicle availability and calcium handling may contribute to the reduced paired-pulse depression observed after aldosterone treatment. Together, these transcriptional changes suggest aldosterone may regulate multiple components of presynaptic release machinery to support increased activity at HSD2 neuron terminals.

Interestingly, the HSD2 neuron response to aldosterone shares features with AgRP “hunger” neurons in response to caloric deficit. Fasting increases *Zbtb16* and *Sv2c* expression in AgRP neurons and similarly induces a broad transcriptional program associated with synaptic function and neurotransmitter release. Importantly, fasting activates the hypothalamic-pituitary-adrenal axis, and glucocorticoid signaling contributes to AgRP neuron function and *Zbtb16* expression^81,92,93^. However, whether *Zbtb16 contributes to the regulation of* synapse-related genes is unknown. Together with aldosterone-induced *Zbtb16*, *Sv2c*, and other synapse-associated genes in HSD2 neurons, these findings suggest that corticosteroid signaling may recruit overlapping transcriptional programs in HSD2 and AgRP neurons to support sustained activity and neurotransmission during distinct physiological deficiency states.

In summary, we demonstrate that MR signaling is essential for HSD2 neuron function and define a molecular signature of aldosterone-MR signaling in neurons. MR is essential in HSD2 neurons for sodium appetite in response to aldosterone treatment and sodium deficiency and also establishes or maintains HSD2 neuron identity or survival during development. Aldosterone-MR signaling increases HSD2 neuron activity while inducing transcriptional and functional adaptations in presynaptic transmission to support sustained neurotransmitter release during elevated firing. Furthermore, the identification of SV2C as an aldosterone-MR-regulated mediator of sodium appetite in HSD2 neurons provides a functional link between aldosterone-responsive gene expression and sodium intake. These findings reveal a neuromolecular framework through which aldosterone-MR signaling regulates HSD2 neuron activity and synaptic function to promote sodium appetite.

## Resource availability

### Lead contact

Further information and requests for resources and reagents should be directed to and will be fulfilled by the lead contact, Jon Resch.

### Materials availability

All unique reagents generated in this study are available from the lead contact with a completed materials transfer agreement.

### Data and code availability

Mouse snRNA-seq data from this study have been deposited into the NCBI Gene Expression Omnibus (GEO) with accession number: GSE326616. Smart-seq2 scRNA-seq of HSD2 neurons following sodium deficiency are available downloading the publicly available dataset at GEO: GSE102332^14^. All original code and Seurat objects have been deposited to Zenodo: https://doi.org/10.5281/zenodo.19343900. Finally, any additional information required to reanalyze the data reported in this paper is available from the lead contact upon request.

## Acknowledgements

We would like to thank members of the Resch and Geerling laboratories for helpful discussions; Jean-Charles Paterna, Lazaros Vasilikos, and the Viral Vector Facility of the Neuroscience Center Zurich (ZNZ) at University of Zurich for helpful discussions and technical assistance with cloning and AAV production; Kevin Knudtson, Garry Hauser, Mary Boes, Einat Snir, and the University of Iowa Carver College of Medicine Iowa Institute for Human Genetics Genomics Division. (IIHG, RRID: SCR_023422) for helpful discussions and technical assistance with single-nucleus RNA sequencing; Linus Tsai, Christopher Jacobs, Anton Gulko, and the BNORC Functional Genomics and Bioinformatics Core (P30DK046200) for helpful discussions and technical assistance; Heath Vignes, Michael Shey, Thomas Kaufman, and the University of Iowa Carver College of Medicine Flow Cytometry Facility for helpful discussions and technical assistance. This research was funded by the following NIH grants to J.M.R.: R00HL144923 and R01NS141072, and to M.C.M.: F31HL170784. T.C.B. and M.C.M. were also supported by an NIH predoctoral training grant in the pharmacological sciences (T32GM144636). Funding from the American Heart Association (AHA) was awarded to J.M.R. for a Career Development Award (AHA 935362), and A.K. was supported by an AHA Diversity Supplement (AHA 24DIVSUP1285051). Additional support came from the University of Iowa Fraternal Order of Eagles Diabetes Research Center Pilot and Feasibility Catalyst Grant, a Brain Research Foundation Seed Grant (BRFSG-2024-12), and an Iowa Neuroscience Institute Early Stage Investigator award from the Carver Charitable Trust.

## Author Contributions

Conceptualization, M.C.M., A.K., and J.M.R.; data curation, M.C.M., A.K., T.C.B., M.R.S., and A.B.B.; formal analysis, M.C.M., A.K., and T.C.B.; funding acquisition, M.C.M., A.K., and J.M.R.; investigation, M.C.M (behavioral experiments, FISH/IF); M.C.M. and A.K. (CRISPR guide design and cloning); M.C.M. and T.C.B. (snRNA-seq); A.K. (electrophysiology); methodology, M.C.M., A.K., S.G., J.C.G., and J.M.R.; project administration, J.M.R.; resources, J.M.R.; software, M.C.M. and T.C.B.; supervision, M.C.M., A.K., and J.M.R.; validation, M.C.M., A.K., T.C.B., and J.M.R.; visualization, M.C.M. and J.M.R.; writing – original draft, M.C.M. and J.M.R.; writing – review and editing, M.C.M., A.K., and J.M.R. All authors reviewed and approved the final version of the manuscript.

## Declaration of generative AI and AI-assisted technologies in the writing process

During the preparation of this work, the author(s) used ChatGPT in order to improve the clarity and readability of the text. After using this tool/service, the author(s) reviewed and edited the content as needed and take(s) full responsibility for the content of the publication.

## Methods

### Mice

All animal care and experiments were approved by the Institute for Animal Care and Use Committee (IACUC) at the University of Iowa (3102343). Prior to starting experiments, all mice were housed with a 12-hour light-dark cycle and had *ad libitum* access to water and a standard chow diet (Inotiv 7913). Male and female mice aged 5-14 weeks were used for hormone measurement, snRNA-seq, and electrophysiology studies. Male mice were used for behavioral experiments, as female rodents display reduced sodium appetite^15,94,95^. Nevertheless, this limits our ability to extend our sodium appetite behavior findings to females. No sex-dependent effects were observed in snRNA sequencing analyses. *Hsd11b2*-IRES-Cre (JAX:030545)^15^, *Nr3c2*^lox/lox^ ^33^, *Nestin*-Cre (JAX:003771)^34^, *Syn1*-Cre (JAX:003966)^35^, and *Slc17a6*-IRES-Cre (JAX:028863)^96^ mice were maintained on a mixed background, and all experimental animals were hemizygous or heterozygous for genetic modifications except *Nr3c2*^lox/lox^ mice, which were homozygous.

### Cre-dependent AAV-CRISPR-SaCas9 design and construction

Cre-dependent CRISPR-SaCas9 AAVs targeting *Rosa26, Nr3c2,* and *Sv2c* were designed as previously described by Hunker et al., 2020, including the use of the same *Rosa26* guide RNA^37^. The *Nr3c2* and *Sv2c* RNA guides were designed using the BROAD Institute’s CRISPick tool^97,98^ and the CRISPOR design tool^99^. The CRISPick tool utilizes Azimuth 2.0 to calculate “on-target” efficiency, based on Doench et al., 2016^97^. The CRISPOR design tool uses Hsu et al., 2013^100^ and Tycko et al., 2018^101^ to predict “off-target” cut sites and calculate a specificity score, and Bae et al., 2014^102^ to predict the percentage of mutations leading to frame shifts, “out-of-frame.” The *Nr3c2* guide (5’ GGCCTAGATATGGAAAGGCGC 3’) targeted the first coding exon; BROAD efficiency score, 0.8623; CRISPOR specificity score, 90; out-of-frame, 66%. Predicted off-target editing sites are listed in **Table S3A**. The *Sv2c* guide (5’ CTGGAACCGACTGTAGGACCG 3’) targeted the first coding exon; BROAD efficiency score, 0.6628; CRISPOR specificity score, 100; out-of-frame, 57%. Predicted off-target editing sites are listed in **Table S3B**. Oligonucleotides were ordered (Integrated DNA Technologies) for *Rosa26* and *Nr3c2* sgRNAs with a 5’ CACCG-3’ overhang on the forward primer and a 5’ AAAC--C 3’ overhang on the reverse primer to allow for integration into a pAAV-DIO-SaCas9-HA-U6-sgRNA plasmid (Addgene, 124844)^37^. Primers used: *Rosa26* forward - 5’ CACCGCTCGATGGAAAATACTCCGAG 3’, *Rosa26* reverse - 5’ AAACCTCGGAGTATTTTCCATCGAGC 3’, *Nr3c2* forward - 5’ CACCGGGCCTAGATATGGAAAGGCGC 3’, *Nr3c2* reverse - 5’ AAACGCGCCTTTCCATATCTAGGCCC 3’. For ligation, pAAV-DIO-SaCas9-HA-U6-sgRNA was digested with BsaI-HF (NEB, R3535) overnight at 37°C. Following digestion, the plasmid backbone was purified using a QIAGEN Miniprep Kit (QIAGEN, 27104), according to the manufacturer’s instructions. Oligos were phosphorylated using T4 Polynucleotide Kinase (NEB, M0201) and annealed into the plasmid backbone using the T4 DNA Ligase (NEB, M0202). The ligated plasmid was transformed into Stbl3 chemically competent *E. coli* cells (Invitrogen, C737303) and plated on LB-agar plates containing ampicillin (Sigma-Aldrich, A9518-5G). After overnight incubation at 37°C, colonies were screened by colony PCR using primers flanking the sgRNA insertion site. Positive clones were grown in LB containing ampicillin overnight and subjected to plasmid miniprep (QIAGEN, 27104). Plasmid inserts were confirmed via Sanger sequencing. A single validated clone was then expanded in a larger culture and processed using a Maxiprep kit (Macherey-Nagel, 740424.10). We again validated the purified plasmid by Sanger sequencing and SmaI digestion to confirm ITR integrity (Inverted Terminal Repeats). The final validated plasmid was then submitted to the Viral Vector Facility of the Neuroscience Center Zurich (ZNZ) for viral packaging with an AAV9 serotype. The *Sv2c* guide RNA was similarly cloned into the pAAV-DIO-SaCas9-HA-U6-sgRNA backbone, sequence validated, and packaged with an AAV9 serotype by the Viral Vector Facility of the ZNZ.

### Cre-dependent AAV-shRNA design and construction

Cre-dependent shRNA targeting *Sv2c* was designed, validated, and packaged with an AAV9 serotype by Vector Biolabs (AAV9-CreON-(EGFP-U6-m-SV2C-shRNA)). In brief, candidate shRNA target sequences were designed based on computational algorithms, screened for sequence similarities to other genes, and transfected into HEK293 reporter cell lines for knockdown of *Sv2c*. The chosen sh*Sv2c* (5’ GCAGATTCCTCGGAGTCAAGT 3’) achieved 85% knockdown of *Sv2c* mRNA, as assessed by a dual-luciferase assay. The sh*Sv2c* design consisted of the target sequence (GCAGATTCCTCGGAGTCAAGT), a hairpin loop sequence (CTCGAG), and a termination sequence (TTTTT). As a result, the specific sequence inserted into pAAV-CreON-EGFP-U6-m-SV2C-shRNA was: 5’- ACC GCAGATTCCTCGGAGTCAAGT CTCGAG ACTTGACTCC GAGGAATCTG C TTTTT-3’. The control virus contained a scrambled target sequence (TTGGTGCTCTTCATCTTGTTG); AAV9-CreON-(EGFP-U6-shScrmb-SHC002); Vector Biolabs, 7045).

### Surgical procedures

#### Stereotaxic injections

Mice were anesthetized with a ketamine/xylazine cocktail (100 mg/kg and 10 mg/kg, respectively) prepared in 0.9% sterile saline. To target the nucleus of the solitary tract (NTS), mice were placed into the ear bars of a stereotaxic frame (Kopf, 942-A) with the nose angled down at ∼45°. A midline incision was then made on the back of the skull, and the skin and muscles were retracted to expose the dura mater covering the fourth ventricle at the cisterna magna. A portion of the dura was then excised using a 29-gauge needle to allow access to the brainstem. Bilateral injections of adeno-associated virus (AAV; 200nL/side) were made using a pulled glass micropipette (20-40 μm diameter tip) expelled by picoliter air puffs through a solenoid valve controlled by a Grass stimulator (∼10 nL/second). The following coordinates were used, in reference to calamus scriptorius: anterior +0.3 mm, lateral ± 0.15 mm, and ventral −0.3 mm. The pipette was withdrawn 1 minute post-injection to minimize backflow. After completing the injections, we closed the incision first with absorbable suture on the muscle layer (MedVet International, JORG22419), then with non-absorbable suture for the skin (MedVet International, MV-8685-V), and finally with topical tissue adhesive (Vetbond, 3M). Mice were placed on a heated pad for recovery and given subcutaneous injections of Meloxicam (4 mg/kg) for 48 hours for post-op analgesia. Mice were allowed a minimum of 10 days of recovery post-stereotaxic surgery before acclimation to experimental conditions.

The following AAV-SaCas9 viruses were custom-packaged by the Viral Vector Facility of the Neuroscience Center Zurich (ZNZ) and used to induce indel mutations and for subsequent sodium appetite behavioral assays and electrophysiology: AAV9-CMV-DIO-SaCas9-HA-U6-sg*Rosa26*^37^, AAV9-CMV-DIO-SaCas9-HA-U6-sg*Nr3c2*, and AAV9-CMV-DIO-SaCas9-HA-U6-sg*Sv2c*-HA. For shRNA-mediated knockdown of SV2C for sodium appetite behavioral assays, we injected AAV9-CreON-(EGFP-U6-m-SV2C-shRNA) (Vector Biolabs, shAAV-273493) and AAV9-CreON-(EGFP-U6-shScrmb-SHC002) (Vector Biolabs, 7045) as a control. For overexpression of Sv2c for sodium appetite assessment, we injected AAV9-CMV-DIO-mSV2C-P2A-GFP (Vector Biolabs, 273493) or AAV8-hSyn-DIO-GFP (Addgene, 50457) as a control. AAV8-hSyn-DIO-mCherry (Addgene, 50459) was used for a control in sodium appetite behavioral assays and for cell-attached recordings of HSD2 neuron firing rate alone or mixed 1:1 with either AAV9-CMV-DIO-SaCas9-HA-U6-sg*Rosa26*-HA or AAV9-CMV-DIO-SaCas9-HA-U6-sg*Nr3c2*-HA. HSD2 neuron terminal stimulation electrophysiology was performed with AAV5-Syn-DIO-rc[ChrimsonR-tdTomato] (Addgene, 62723). For fluorescence-activated nuclei sorting (FANS), AAVDJ-hSyn-DIO-H2B mCherry^42^ was custom packaged by the Viral Vector Facility of the Neuroscience Center Zurich (ZNZ) and injected alone or mixed 1:30 with either AAV9-CMV-DIO-SaCas9-HA-U6-sg*Rosa26*-HA or AAV9-CMV-DIO-SaCas9-HA-U6-sg*Nr3c2*-HA to assess editing efficiency of AAV-Cas9 vectors.

#### Osmotic pump implantation

For peripheral infusion of aldosterone, osmotic pumps (Alzet model 1007D or RWD model 1001W) were primed with aldosterone or vehicle the afternoon before implantation. Mice were anesthetized using isoflurane at a rate of 5% for induction and 0.3% to maintain a surgical plane. The pump was implanted subcutaneously above the scapular region and the incision was closed with sutures followed by topical application of tissue adhesive.

### CRISPR-Cas9 on-target indel mutation confirmation

Targeted sequencing of amplicons from Cre-dependent AAV-CRISPR-Cas9 injected mice was performed as previously published^37^, with minor modifications. For isolation of AAV-CRISPR-Cas9-expressing HSD2 neurons, AAVDJ-hSyn-DIO-H2B-mCherry was co-injected with AAV9-CMV-DIO-SaCas9-sg*Rosa26* or - sg*Nr3c2* at a 1:30 ratio. 21 days following surgery, brains were extracted and tissue prepared as described below, without the use of RNA-protect. 25 mCherry-positive nuclei were sorted using a 100 μm nozzle directly into Armadillo 96-well PCR plates (Thermofisher, AB2396) containing REPLI-g Advanced Single Cell storage buffer. Immediately, whole genome amplification (WGA) using the REPLI-g Advanced Single Cell kit (QIAGEN, 150365) was performed according to the manufacturer’s protocol. Then, the region surrounding the guide target sequence was amplified by two rounds of PCR to increase final DNA yield and target amplification specificity. WGA DNA was diluted 1:50 and amplified using Titanium® Taq DNA Polymerase (Clontech Takara, 639242). That product was then diluted 1:100 and amplified again, using primers with the addition of partial Illumina adaptors. PCR 1 protocol: Initial denaturation at 95°C for 5 minutes. 35 cycles of denaturing at 95°C for 30 seconds, annealing at 58°C for 30 seconds, and extension at 72°C for 30 seconds, followed by a final extension at 72°C for 10 minutes. PCR 2 protocol was identical except the annealing occurred at 65°C. A previously published primer set for *Rosa26* was used^37^: primer set 1 forward: AAAGTCGCTCTGAGTTGTTATCAG; primer set 2 forward: AAAGGCTAACCTGGTGTGTGG; primer set 1 and 2 reverse: GGAGCGGGAGAAATGGATATG. For *Nr3c2*: primer set 1 and 2 forward: GGAAACCAAAGGCTACCACA; primer set 1 reverse: CAGTTTCCCAGTGCACAGTC; primer set 2 reverse: CAAAACATGGAGCAGCTGGT. Following the second amplification, PCR products were purified by gel extractions using MinElute gel extraction kit (QIAGEN, 28004) and stored at 20°C. Gel-extracted samples were then sent to Genewiz/Azenta for ‘Amplicon EZ’ targeted sequencing of amplicons. From the sequenced regions, we extracted the 4 base pairs upstream of the PAM and 10 base pairs downstream of the guide RNA binding sequence. We then analyzed the proportion of reads that were “wild-type” or contained base-pair changes, insertions, or deletions.

### Sodium appetite behavioral testing

Sodium appetite assays were performed in BioDAQ food/liquid intake monitoring cages. Mice were allowed to recover at least 10 days post-AAV injection (if performed) before being transferred to BioDAQ cages where they were single-housed and had *ad libitum* access to standard chow diet, water, and 0.5 M NaCl. After acclimation, mice underwent dietary sodium deprivation, sodium depletion, and/or aldosterone infusion.

#### Dietary sodium deprivation

Mice were provided clean cages with *ad libitum* access to water and Na^+^-deficient diet without 0.5 M NaCl access for 7 days (Teklad/Envigo, TD-130591). Fresh cages were provided daily for the first 3 days of the 7-day deprivation protocol. Non-sodium-deprived control mice were single-housed with no access to saline, but *ad libitum* access to water and a standard chow diet. Fresh cages were provided daily for the first 3 days as in the deprivation protocol. After 7 days, mice underwent sodium appetite behavioral testing where access to 0.5 M NaCl was returned 30 minutes before the onset of the dark cycle, and intake of 0.5 M NaCl and water was reported for four hours.

Additional animals underwent this protocol and were sacrificed without saline access for plasma aldosterone measurements or assessment of c-Fos in HSD2 neurons.

#### Sodium Depletion

Mice were injected with furosemide (50 mg/kg IP), then provided clean cages with *ad libitum* access to water and either Na^+^ deficient diet or standard chow diet (control) without 0.5 M NaCl access. The following day, mice underwent sodium appetite behavioral testing where 0.5 M NaCl was returned 30 minutes before the onset of the dark cycle, and intake of 0.5 M NaCl and water was reported for four hours. Additional animals underwent this protocol and were sacrificed without saline access for plasma aldosterone measurements.

#### Aldosterone infusion

Intake of 0.5M NaCl and water was continuously monitored for three days prior to implantation of aldosterone-filled osmotic pumps (Alzet model 1007D or RWD model 1001W). Osmotic pumps were filled with either aldosterone (0.5 mg/ml aldosterone with 2.5% EtOH in 0.9% saline) or vehicle (2.5% EtOH in 0.9% saline). The rate of aldosterone release was 250ng/hr. After pump implantation, mice underwent sodium appetite behavioral testing where intake of 0.5M NaCl and water were recorded for seven days. Due to a miscalculation of the aldosterone dose for HSD2^Cas9-sg*Rosa*26^/HSD2^Cas9-sg*Sv2c*^ mice and one cohort of HSD2^shScrmbl^/HSD2^sh*Sv2c*^ mice, these animals received approximately twice the intended aldosterone dose during aldosterone-induced sodium appetite behavioral testing, resulting in higher saline intake than in other studies. For assessment of c-Fos in HSD2 neurons, mice were perfused upon removal from BioDAQ cages at the end of the seventh day. Additional animals underwent this protocol and were sacrificed after 24 hours for c-Fos immunofluorescence and electrophysiological recordings of firing rate from HSD2 neurons, or after seven days for plasma aldosterone measurements.

### Plasma aldosterone measurements

To measure plasma aldosterone levels, we performed an Enzyme Linked ImmunoSorbent Assay (ELISA) (IBL, IB79134). Trunk blood was collected in K2 and EDTA coated tubes (BD, 367841) between 8-10 am following rapid decapitation of mice. Blood was collected following dietary sodium deprivation, sodium depletion via furosemide, or 7 days of aldosterone infusion. The blood was then centrifugated at 1,800 rcf for 15 mins at 4°C. Subsequently, plasma was collected and aliquots were stored at −80°C. All plasma was diluted 1:10 in plasma diluent for the aldosterone ELISA, whereafter the standard instructions for the kit were followed.

### Single-nucleus RNA Sequencing

#### Tissue collection, library preparation, and sequencing

We performed HSD2-neuron-enriched snRNA-seq on the dorsal vagal complex (DVC) from 3 replicates of vehicle- and aldosterone-treated mice (n = ∼9 mice per sex/treatment). *Hsd11b2*-IRES-Cre mice aged 5-10 weeks received bilateral stereotaxic injections of Cre-dependent AAV-H2B-mCherry into the NTS. After 13 days of recovery, mice were implanted with vehicle- or aldosterone-filled osmotic pumps between 9:00 am – 12:00 pm. The following morning, mice were sacrificed between 9:00 am – 12:00 pm by rapid decapitation immediately after removal from the home cage. Brains were extracted and chilled in DMEM/F12 media slush, then placed dorsal side up in a chilled stainless steel brain matrix (Roboz Surgical Instrument Co., SA-2175), and 1 mm coronal sections of the caudal medulla were collected.

For snRNA-seq using 10X, 1 mm coronal slices from the caudal medulla were placed in chilled RNAprotect® solution (QIAGEN, 76106), and then the caudal NTS and surrounding area was microdissected using a 1 mm diameter punch under a fluorescent stereoscope (Nikon SMZ-800N Stereoscope). During dissections, tissue was pooled according to sex and treatment (vehicle or aldosterone). All pooled samples were incubated overnight at 4°C in RNAprotect®. RNAprotect® solution was then removed from sample tubes the following day before storing samples at −70°C. Each sample underwent nuclear dissociation and purification procedures before being incubated with NucBlue (Thermo Fisher Scientific), as previously described^103,104^. After incubation, nuclei were sorted into RT Reagent B (10x Genomics) using a Cytek Aurora CS cell sorter with a 70 μm nozzle to remove poor quality nuclei and enrich for HSD2 neurons. First, 4,000 unbiased DVC nuclei were collected, followed by collection of all remaining mCherry^+^ nuclei (HSD2 neurons) in the sample, ranging from 278 to 715. Sorted nuclei were immediately loaded onto the 10x Chromium controller (10x Genomics). cDNA and gene expression libraries were generated according to the manufacturer’s instructions (10x Genomics, 3’ GEM-X chemistry). Gene expression libraries were sequenced on either an Illumina NovaSeq 6000 or on an AVITI24 (Element Biosciences) with a targeted depth of 100,000 reads per nucleus. One replicate was performed to capture single-nucleus multiome ATAC + Gene Expression. Tissue from these samples was frozen on dry ice instead of treating with RNAprotect®. After nuclei dissociation and purification as above, 4 µl of resuspended nuclei at ∼1,000 nuclei/µl were added to a collection tube with ATAC Buffer B (10X Genomics). Subsequently, mCherry+ nuclei (475/group) were flow-sorted into the same collection tube using a FACS Aria cell sorter with a 70 μm nozzle. Gene expression and ATAC libraries were sequenced on an Illumina NovaSeq 6000. 10X Genomics Cell Ranger was used to map reads to the GRCm38 reference genome and generate feature-barcode matrices.

#### Single-nucleus RNA sequencing quality assessment, data integration, and analysis

Data from 10X Chromium-v4 and 10X Multiome were loaded into an RStudio environment (R v4.4.1) and processed through a custom Seurat-based analysis pipeline run in Seurat v5. We removed nuclei with fewer than 250 unique genes, fewer than 1,000 total UMI, or greater than 5% of mitochondrial gene expression (quantified using *PercentageFeatureSet()*). After filtering, 3 batches of sequencing runs were merged into a single Seurat object (10X Chromium-v4 = 2 batches and 10X Multiome = 1 batch). All “RNA” assay layers were split by replicate, raw counts were log-normalized using Seurat *NormalizeData()*, and the top 5,000 highly variable genes were selected with *FindVariableFeatures()*. The data were then scaled using *ScaleData()*, with percent mitochondrial gene expression regressed, before principal component analysis was performed using *RunPCA()*. Next, *IntegrateLayers()* was used to perform a reciprocal principal component analysis (RPCA)-based integration, generating a batch-effect corrected dimensional reduction titled “integrated.rpca”. Following integration, the top 30 integrated.rpca dimensions were used to construct a nearest-neighbor graph with *FindNeighbors()* before clusters were identified with *FindClusters()*. After identifying clusters, *RunUMAP()* was used to generate a final Uniform Manifold Approximation and Projection (UMAP) dimensional reduction titled “umap.rpca”. Prior to performing differential gene expression analysis (DGEA), RNA-assay layers were re-joined using *JoinLayers()*, and DGEA was completed with *FindAllMarkers()* utilizing the non-parametric Wilcoxon Rank-Sum test with thresholds of > 0.2 average log_2_ fold change and < 0.01 Bonferroni-corrected p-value. These marker genes allowed for identification of doublets/multiplets (clusters containing high expression of cell type markers denoting two or more separate cell types, i.e. oligodendrocytes and macrophages) and low-quality clusters (clusters with high expression of mitochondrial genes or lacking in strong cell-type identifying markers). Once this process was completed for all nuclei within our dataset, we subset for only neuronal nuclei (*Rbfox3*^+^ and *Tubb3*^+^) and repeated the analysis. At this stage, we subset and repeated the analysis for three neuronal subtypes: GABAergic neurons (*Slc321a1*^+^), cholinergic neurons (*Chat*^+^), and glutamatergic neurons (*Slc17a6*^+^). We then identified genes differentially expressed by aldosterone treatment within each individual cluster for all neuronal subtypes. For analysis, we used Seurat’s *FindMarkers()*, utilizing DESeq2 with an average log_2_ fold change threshold of > 0.25 and a Bonferroni-corrected p-value threshold of < 0.05.

#### Gene Ontology analysis

Gene Ontology (GO) enrichment analysis was performed on aldosterone-induced upregulated genes in the GLUT1.Hsd11b2 cluster using the clusterProfiler package. Specifically, *enrichGO()*^105^was used to assess enrichment for all three GO categories within the gene list with a Benjamini-Hochberg-corrected p-value cutoff of < 0.05 and a q-value cutoff of < 0.05. Then, the GOSemSim package^106^ and *simplify()* function was used to combine GO terms with semantic similarity using a reduction value of 0.5. Then, to isolate synapse-related genes, the list of GO terms was further filtered to only include those with key words ‘synapse’, ‘synaptic’, ‘vesicle’, ‘neurotransmitter’, ‘exocytosis’, and ‘action potential’.

### Electrophysiology

#### Cell-attached intrinsic firing recordings

For intrinsic firing of HSD2 neurons, 5–8-week-old *Hsd11b2*-IRES-Cre mice received NTS injections of AAV8-hSyn-DIO-mCherry alone or mixed with either AAV9-CMV-DIO-Cas9-sg*Rosa26* or AAV9-CMV-DIO-Cas9-sg*Nr3c2*. Aldosterone-filled osmotic pumps were implanted either 24 hours before recordings from AAV8-hSyn-DIO-mCherry injected mice or 6-8 days before recordings from mixed mCherry/Cas9 AAV injections. 3 weeks post-AAV injection, mice were deeply anesthetized with isofluorane and transcardially perfused with ice-cold sucrose-based cutting solution (in mM: 190 sucrose, 1.9 KCl, 1.2 NaH₂PO₄, 25 NaHCO₃, 0.25 CaCl₂, 6 MgCl₂·6H₂O, 10 dextrose, 3 myo-inositol, 2 L-ascorbate, and 2 sodium pyruvate, continuously bubbled with 95% O₂ and 5% CO₂ (carbogen)). 300 μm coronal brainstem slices containing the nucleus of the solitary tract (NTS) were prepared with a vibratome (Campden 7000smz-2). Slices recovered in artificial cerebrospinal fluid (aCSF, in mM: 124 NaCl, 2.5 KCl, 1.2 NaH₂PO₄, 21.4 NaHCO₃, 2.4 CaCl₂, 1.2 MgCl₂·6H₂O, 10 dextrose) for 30 minutes at 28°C followed by at least 30 minutes at room temperature (20–22°C) before recordings. mCherry fluorescence was used to visualize HSD2 neurons for loose-seal cell-attached recordings^14,107^. Patch pipettes (5-7 MΩ) were filled with aCSF. Recordings were done in the presence of 1mM kynurenic acid and 100 μM picrotoxin to block glutamatergic and GABAergic transmission. Clampex software, a Digidata 1550B digitizer, and MultiClamp 700B amplifier were used to make recordings. Signals were acquired at 10 or 50 kHz, then low-pass filtered at 2 or 10 kHz. Analyses were performed offline using Clampex 1104.

#### Whole-cell voltage-clamp optogenetic paired-pulse recordings

For measuring evoked excitatory postsynaptic currents from paired-pulse ratio experiments, 5–8-week-old *Hsd11b2*-IRES-Cre mice received NTS injections of AAV5-Syn-DIO-rc[ChrimsonR-tdTomato]. Vehicle- or aldosterone-filled osmotic pumps were then implanted 6-8 days before recordings. At least 3 weeks post-AAV injection, 300 μm coronal brain slices containing the BSTvL were prepared and recordings were performed as above with the following modifications. Patch pipettes (3–4 MΩ) were filled with a Cs-methanesulfonate-based internal solution for EPSC recordings (in mM: 135 CH₃O₃SCs,10 HEPES, 1 EGTA, 0.2 CaCl₂, 4 Mg-ATP, 0.3 Na-GTP, 8 Na₂-phosphocreatine, and 8 QX-314; pH adjusted to 7.3 with CsOH, and osmolarity to ∼295 mOsm). Recordings were performed in the presence of tetrodotoxin (TTX; 1 μM), and 4-aminopyridine (4-AP; 500 μM) to isolate monosynaptic EPSCs. BSTvL cells were patched inside the tdTomato-visualized projection field of HSD2 neurons. The protocol for photostimulation of ChrimsonR-expressing fibers comprised of recording sweeps with two 5 ms pulses of 595 nm light from an LED light source, spaced 50 ms apart. Light output was controlled via pulses from the amplifier to the LED. Responses were recorded for 2-15 sweeps. Connectivity rate was calculated as the fraction of patched BSTvL neurons that received light-evoked input. Latency was calculated as the time between light onset and current onset. Paired-pulse ratio was calculated as the response amplitude to the second light pulse divided by the first. The area under the current trace was measured over the 100 ms following each light stimulus. Decay time was measured in Clampfit as the time for the current to fall from 90% to 10% of its peak amplitude. Cells were excluded from the analyses of decay kinetics if the measurements were classified as “not detectable”. This occurred when the response overlapped with the subsequent stimulus or exhibited multiple peaks which prevented reliable characterization of the decay phase.

### Histology

Mice were deeply anesthetized with a ketamine/xylazine cocktail (150 mg/kg and 15 mg/kg, respectively) prepared in 0.9% sterile saline. They were then transcardially perfused with phosphate-buffered saline (PBS; Boston BioProducts, #BM-220) followed by 10% neutral-buffered formalin (NBF; Fisher, #SF100-20). Brains were then extracted and postfixed overnight in 10% NBF at 4°C. Brains were then switched to 20% sucrose in PBS overnight at 4°C before being sectioned at 30 μm on a freezing microtome and stored in a cryoprotectant solution at −20°C for storage.

#### Immunofluorescence

Tissue sections were removed from cryoprotectant and washed 3 times in PBS before being placed in a blocking solution for 1 hour (3% Normal Donkey Serum (EMD Millipore, #S30-100ML) and 0.4% Triton X-100 in 1X PBS). Primary antibody solutions were prepared in blocking solution at the following concentrations for overnight incubation: rabbit anti-HSD2 (Proteintech, 14192-1-AP) at 1:3,000, guinea pig anti-cFos (Synaptic Systems, 226 308) at 1:10,000, and rabbit anti-HA (Cell Signaling Technology, 3724S) at 1:3,000. The next day, tissue sections were washed 4 times with PBS, then placed in a secondary antibody solution prepared in blocking solution for two hours. Alexa Fluor fluorescent secondary antibodies (Thermo Fisher Scientific) at 1:1000 or Cyanine-conjugated fluorescent antibodies (Jackson ImmunoResearch) at 1:1000 were used. All slides were coverslipped using VECTASHIELD Plus mounting media with DAPI (Vector Laboratories, H-2000-2) and scanned at 10X or 20X + Z-stack magnification on an Olympus Slideview VS200 slide-scanning microscope.

#### Immunofluorescence and Opal-TSA multiplexing (sequential labeling of primary antibodies from the same host)

Brain tissue sections were removed from cryoprotectant and washed three times with PBS. Tissue sections were then mounted onto Superfrost Plus microscope slides (Fisher Scientific, 12-550-15) and allowed to dry for 3-6 hours at room temperature. Slides were then washed three times in PBS before incubation for 15 mins in freshly prepared 1% H_2_O_2_ in PBS. Following four additional washes in PBS, the slides were then incubated overnight at room temperature with rabbit anti-HA (Cell Signaling Technology, 3724S) diluted 1:10,000 in blocking solution. The following morning, slides were washed five times in PBS and incubated for 1 hour with a biotinylated horse anti-rabbit secondary antibody (Vector Labs, BA-1100) diluted 1:600 in blocking solution. The slides were then washed five times in PBS for 10 mins each. During this time, ABC solution was prepared: 45 μL Reagent A and 45 μL of Reagent B (Vector Labs, PK-6100) in 10mLs of 1X PBS + 0.4% Triton X-100. Following the washes and 30 mins after the preparation of ABC solution, slides were incubated in ABC solution at room temperature for one hour. Next, the slides were washed three times in PBS followed by three washes in 1X RNAscope Wash Buffer (Advanced Cell Diagnostics, #320058). Slides were incubated with Opal dye 520 (Akoya Biosciences, P1487001KT) at 1:3,000 in TSA Buffer (Advanced Cell Diagnostics, #322810) for 30 mins followed by three washes in RNAscope Wash Buffer and one wash in PBS. To remove the rabbit anti-HA primary, slides were then placed in 1X RNAscope Target Retrieval (Advanced Cell Diagnostics, #322001) solution and microwaved on HIGH power until boiling, followed by 15 mins at 20% power to keep the solution at boiling temperature without losing excessive liquid to evaporation. Once cooled, slides were washed twice more in PBS before being incubated in primary antibodies diluted in blocking solution overnight at room temperature (rabbit anti-HSD2 at 1:1,000 (Proteintech, 14192-1-AP) and guinea pig anti-cFos (Synaptic Systems, 226 308) at 1:10,000). The next day, slides were washed 4 times in PBS then incubated for 1-2 hours with biotinylated anti-rabbit secondary antibody at 1:500, Alexa Fluor fluorescent secondary antibodies (ThermoFisher) at 1:1000 or Cyanine-conjugated fluorescent antibodies (Jackson ImmunoResearch) at 1:1000 in blocking solution. If biotinylated secondary was used, slides were washed 4 times in PBS before incubation with streptavidin-conjugated Alexa Flour 750 (Invitrogen, S21384) diluted 1:1000 in blocking solution for 1 hour. All slides were washed 5 times with PBS before being coverslipped and imaged as described above.

#### Fluorescence in situ hybridization

To label *Nr3c2* (MR), *Sv2c* (SV2C), *Slc9a3* (Sodium-hydrogen antiporter 3), we performed fluorescence *in situ* hybridization (FISH) using the RNAscope Multiplex Fluorescent Reagent Kit V2 (Advanced Cell Diagnostics, #323100) followed by immunofluorescence (IF) for HSD2 and ChAT, using the manufacturer’s protocol with minor modifications. In brief, mice were perfused as described above using 1X RNAse-free PBS (Invitrogen, 10010-023) and 10% NBF. Brains were extracted and postfixed overnight in 10% NBF and then switched to 10%, 20%, and 30% sucrose in 1X RNAse-free PBS for one night each. The brains were then cut at 30 μm on a freezing microtome and placed into PBST (1X RNAse-free PBS with 0.5% Triton X-100). Slices were then washed once more with PBST before being mounted on Superfrost plus slides. Once dried, slides were placed at −70°C for long-term storage. Upon removal from −70°C, slides were washed with PBS at room temperature, dried for 30 mins at 60°C, and then post-fixed in 10% NBF for 15 mins at 4°C. Slides were then dehydrated in a series of 50%, 70%, and 100% (twice) ethanol at room temperature. Brain sections were then covered with RNAscope Hydrogen Peroxide for 10 mins at room temp and dehydrated once more in 100% ethanol before drying at 60°C for 5 mins. A hydrophobic barrier was drawn around the tissue sections (Vector Laboratories, #H-4000) before RNAscope protease III was applied to the sections for 30 mins at 40°C. All remaining steps were performed at 40°C, unless otherwise specified. Slides were then washed with ultrapure water before probe hybridization with Mm-Nr3c2-E5E6 (Advanced Cell Diagnostics, 456321), Mm-Sv2c-C2 (Advanced Cell Diagnostics, 545001-C2), or Mm-Slc9a3 (Advanced Cell Diagnostics, 430001) for 2 hours at 40°C. Subsequently, slides were incubated with AMP-1 for 30 mins, AMP-2 for 30 mins, and then AMP-3 for 15 mins. Slides were then incubated with HRP-C1 for 15 mins, followed by Opal 570 or Opal 650 (Akoya Biosciences, FP1488001KT or FP1496001KT) diluted 1:750 in TSA buffer (Advanced Cell Diagnostics, 322810) for 30 mins, and finally HRP-blocker for 15 mins. For IF, slides were then incubated overnight at 4°C with rabbit anti-HSD2 at 1:1000 and goat anti-ChAT (EMD Millipore, AB144P) at 1:300 in blocking solution. The following morning, slides were washed 4X with PBS, then incubated with Alexa Fluor fluorescent secondary antibodies (ThermoFisher) at 1:1000 in blocking solution for two hours at room temperature. Slides were then washed with PBS 4X before being coverslipped and imaged at 10X, 20X + virtual Z stacks, or 40X + virtual Z-stacks with an Olympus VS200 slide scanning microscope.

### Cell and puncta counting

To count c-Fos^+^ HSD2 neurons, 10X images of a 1-in-3 series of HSD2-containing NTS sections were manually counted as ‘total HSD2 neurons’ and ‘HSD2 neurons with nuclear Fos immunoreactivity’. To count *Sv2c* puncta within HSD2 neurons, 40X + virtual Z-stack images of the HSD2 field were acquired with an Olympus VS200 slide scanning microscope. Z-stacks were then extracted and saved as a tiff file. ImageJ (v2.1.0) was then used to collapse each z-image into a single projection using ‘maximum intensity’, then each fluorescent channel was merged into a single image. ImageJ’s ‘cell counter’ tool was then used to manually count each distinct FISH puncta per HSD2 neuron.

### Statistical analysis

Statistics for snRNA-seq were performed in R, as described above. All other analyses were conducted using GraphPad Prism (v11.0.1), with the specific statistical tests for each experiment indicated in the figure legends. No statistical methods were used to predetermine sample size, and randomization and/or blinding were not applied for snRNA-seq experiment. For sodium appetite assays with two groups, a two-way repeated-measures ANOVA with Geisser-Greenhouse correction was used to analyze intakes throughout the measurement period. A Welch’s t-test was used to analyze total intakes after 4 hours of access (sodium deprivation and sodium depletion), average intake per day (aldosterone infusion), percent c-Fos^+^ HSD2 neurons following aldosterone treatment, mRNA puncta per HSD2 neuron, and electrophysiology data. For sodium appetite assays with three groups, a one-way ANOVA with Tukey multiple-comparisons correction was used to analyze total intake after 4 hours of 0.5M NaCl access (sodium deprivation and sodium depletion) or average intake per day (aldosterone infusion). For plasma aldosterone measurements and percent c-Fos^+^ HSD2 neurons following dietary sodium deprivation, a one-way Brown-Forsythe and Welch ANOVA tests were performed with Dunnett T3 correction. All results are presented as mean ± SEM, except for electrophysiology and puncta per HSD2 neuron, which were represented as median and quartiles. Statistical significance was defined as an alpha value of less than 0.05.

## Supplemental Items

Table S1. snRNA-seq marker gene lists, related to Figure 3. A) All cell markers. B) Glutamatergic neuron markers. C) GABAergic neuron markers. D) Cholinergic neuron markers.

Table S2. snRNA-seq aldosterone-induced DEGs, related to Figure 3. A) Glutamatergic cluster DEGs. B) GABAergic cluster DEGs. C) Cholinergic cluster DEGs.

Table S3. CRISPR-Cas9 guide predicted off-target editing sites, related to Figure 2 and Figure 5. A) Cas9-sgNr3c2 guide. B) Cas9-sgSv2c guide.

**Figure S1:**
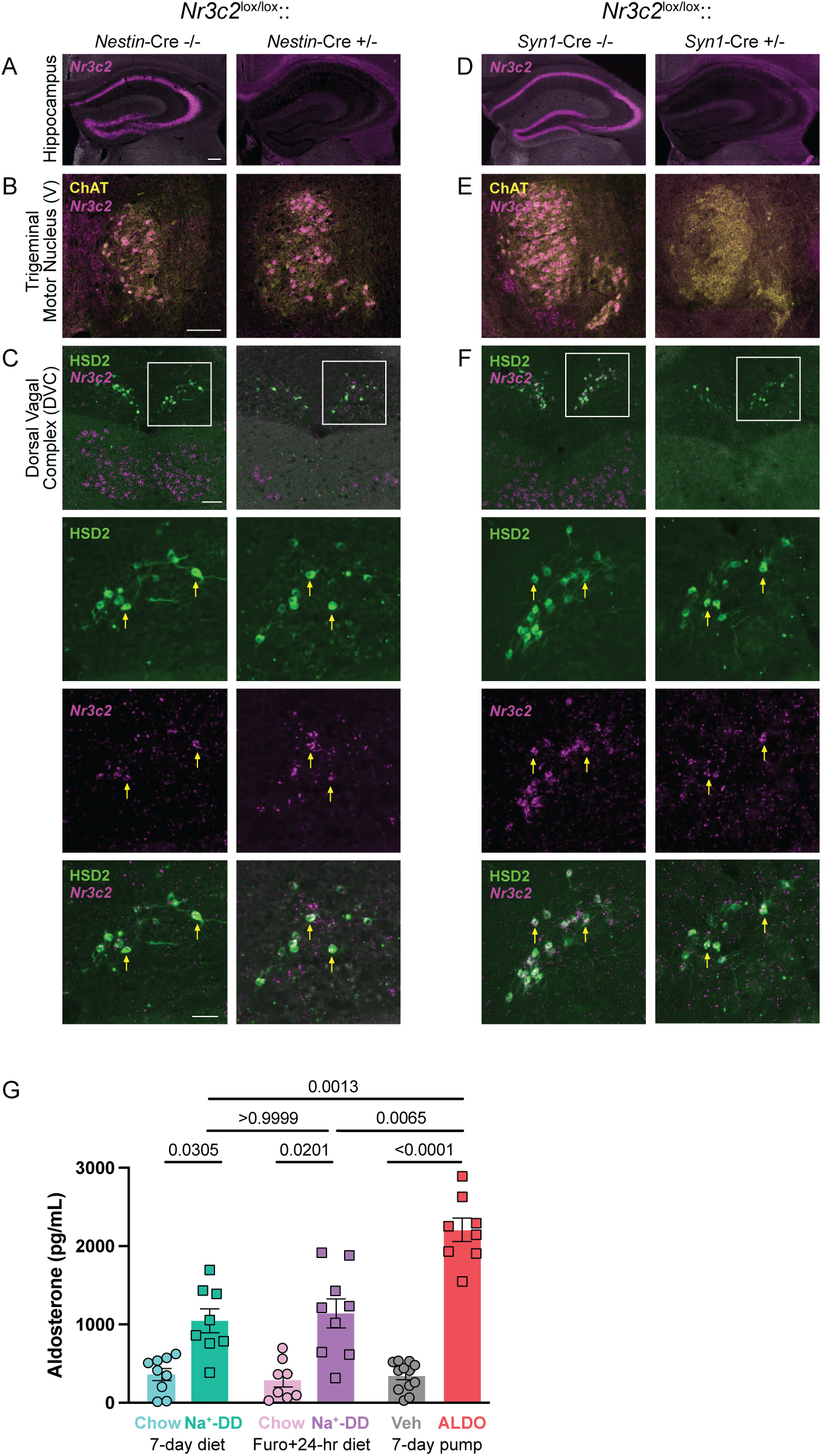
*Nr3c2*^lox/lox^ mice crossed to pan-neuronal Cre drivers cause incomplete knockout of MR from HSD2 neurons, related to Figure 1. **A)** *Nr3c2* fluorescence *in situ* hybridization (FISH) in the hippocampus of *Nr3c2*^lox/lox^ (left) and *Nestin*-Cre::*Nr3c2*^lox/lox^ mice (right; scale bar = 200 μm). **B)** *Nr3c2* FISH with colocalization with ChAT immunofluorescence (IF) in the trigeminal motor nucleus of *Nr3c2*^lox/lox^ (left) and *Nestin*-Cre::*Nr3c2*^lox/lox^ mice (right; scale bar = 100 μm). **C)** Representative brain sections of the dorsal vagal complex from *Nr3c2*^lox/lox^ (left) and *Nestin*-Cre::*Nr3c2*^lox/lox^ mice (right), showing in descending order: low-magnification view of *Nr3c2* FISH and HSD2 IF (scale bar = 100 μm), high-magnification view of HSD2 IF in boxed regions, high-magnification view of *Nr3c2* FISH in boxed regions, and high-magnification view of both HSD2 IF and *Nr3c2* FISH (scale bar = 50 μm). Yellow arrows indicate representative co-labeled neurons. **D)** *Nr3c2* FISH in the hippocampus of *Nr3c2*^lox/lox^ (left) and *Syn1*-Cre::*Nr3c2*^lox/lox^ mice (right; scale bar = 200 μm). **E)** *Nr3c2* FISH with colocalization with ChAT IF in the trigeminal motor nucleus of *Nr3c2*^lox/lox^ (left) and *Syn1*-Cre::*Nr3c2*^lox/lox^ mice (right; scale bar = 100 μm). **F)** Representative brain sections of the dorsal vagal complex from *Nr3c2*^lox/lox^ (left) and *Syn1*-Cre::*Nr3c2*^lox/lox^ mice (right), showing in descending order: low-magnification view of *Nr3c2* FISH and HSD2 IF (scale bar = 100 μm), high-magnification view of HSD2 IF in boxed regions, high-magnification view of *Nr3c2* FISH in boxed regions, and high-magnification view of both HSD2 IF and *Nr3c2* FISH (scale bar = 50 μm). Yellow arrows indicate representative co-labeled neurons. **G)** Plasma aldosterone measurements in response to different sodium appetite models used, including the dietary sodium deprivation sodium appetite model (7 days on chow or Na^+^-deficient diet (Na^+^-DD)), the furosemide (Furo)-induced sodium depletion sodium appetite model (Furo + 24 hours on chow or Na^+^-DD), and the aldosterone (ALDO)-induced sodium appetite model (7 days of osmotic pump infusion with vehicle (Veh) or ALDO). All results are presented as mean ± SEM. Statistical analysis was performed using one-way Brown-Forsythe and Welch ANOVA tests with Dunnett T3 correction. p < 0.05 was considered statistically significant.

**Figure S2:**
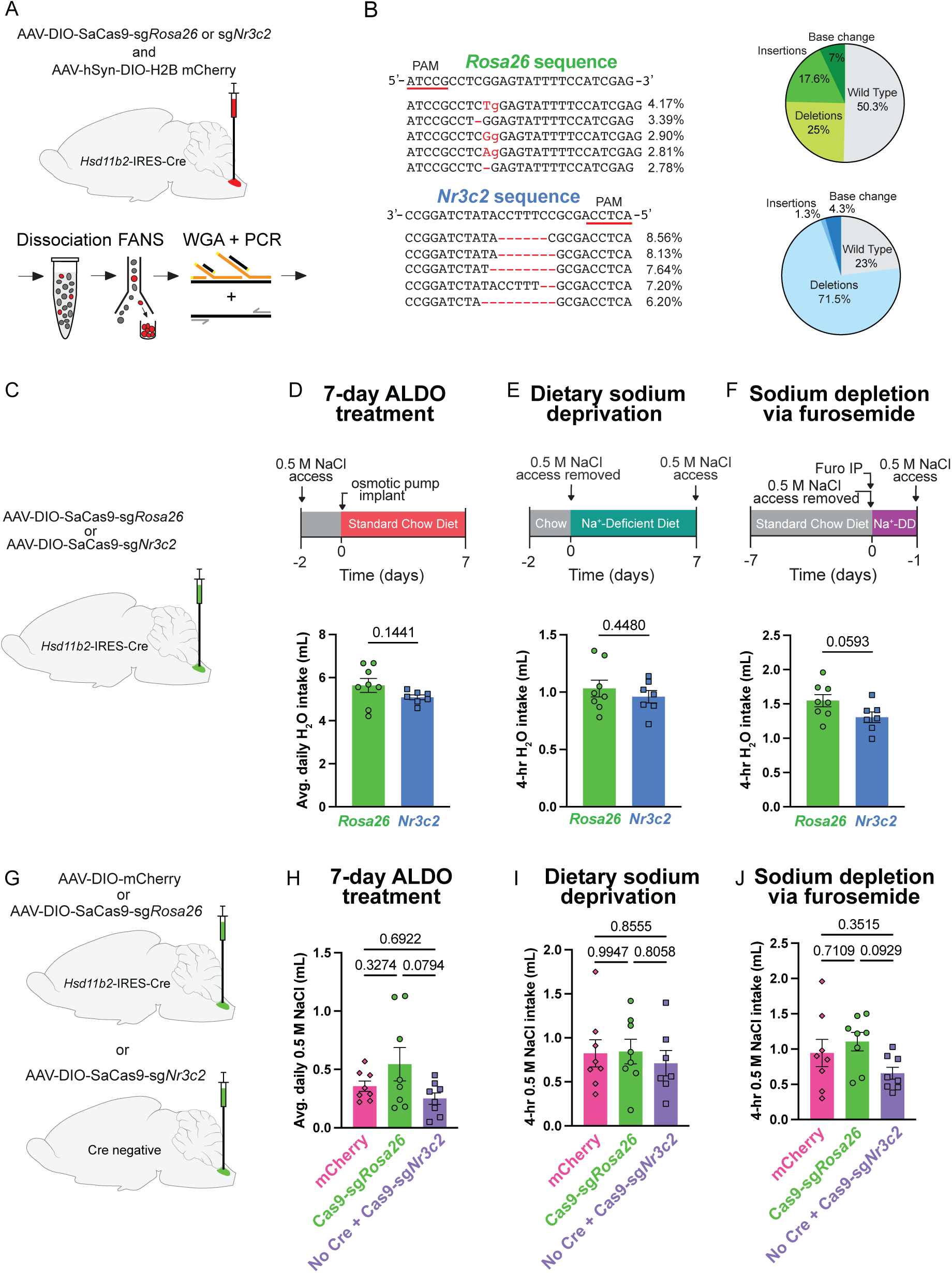
CRISPR-Cas9 AAV strategy effectively disrupts *Nr3c2* in HSD2 neurons, related to Figure 2. **A)** Schematic showing injection of Cre-dependent CRISPR-Cas9 AAV mixed with Cre-dependent AAV-H2B-mCherry into *Hsd11b2*-IRES-Cre mice and subsequent sequencing of Cas9-targeted regions. **B)** The five most commonly detected sequence edits in HSD2^Cas9-sg*Rosa*26^ (top left) and HSD2^Cas9-sg*Nr3c2*^ (bottom left) injected mice, along with the proportion of deletions, insertions, and base pair changes detected for both viruses (right). **C)** Schematic for injection of Cre-dependent CRISPR-Cas9 AAVs into *Hsd11b2*-IRES-Cre mice. **D)** Schematic describing the aldosterone-induced sodium appetite model (top) and average H_2_O intake per day (bottom; n = 7-8 mice/group). ALDO = aldosterone. **E)** Schematic describing the dietary sodium deprivation sodium appetite model (top) and total H_2_O intake in 4 hours (bottom; n = 7-8 mice/group). **F)** Schematic describing the furosemide-induced sodium depletion sodium appetite model (top) and total H_2_O intake in 4 hours (n = 7-8 mice/group). Furo = furosemide; Na^+^-DD = Na^+^-deficient diet. **G)** Schematic for injection of Cre-dependent AAV-Cas9-sg*Rosa26* or AAV-mCherry injection into *Hsd11b2*-IRES-Cre mice (left) and injection of Cre-dependent AAV-Cas9-sg*Nr3c2* into Cre-negative mice (right). **H)** Average daily 0.5 M NaCl intake during aldosterone treatment (n = 7-8 mice/group). **(I)** Total 0.5 M NaCl intake over 4 hours following dietary sodium deprivation (n = 7-8 mice/group) and **(J)** following sodium depletion via furosemide (n = 7-8 mice/group). All 0.5 M NaCl and H_2_O intake data are represented as mean ± SEM. Statistical analysis of saline intake over time between groups was performed using a two-way repeated-measures ANOVA with Geisser-Greenhouse correction. Two-group comparisons without repeated-measures were performed with a Welch’s t-test, and three-group comparisons were performed using a one-way ANOVA with Tukey multiple-comparisons correction. p < 0.05 was considered statistically significant.

**Figure S3:**
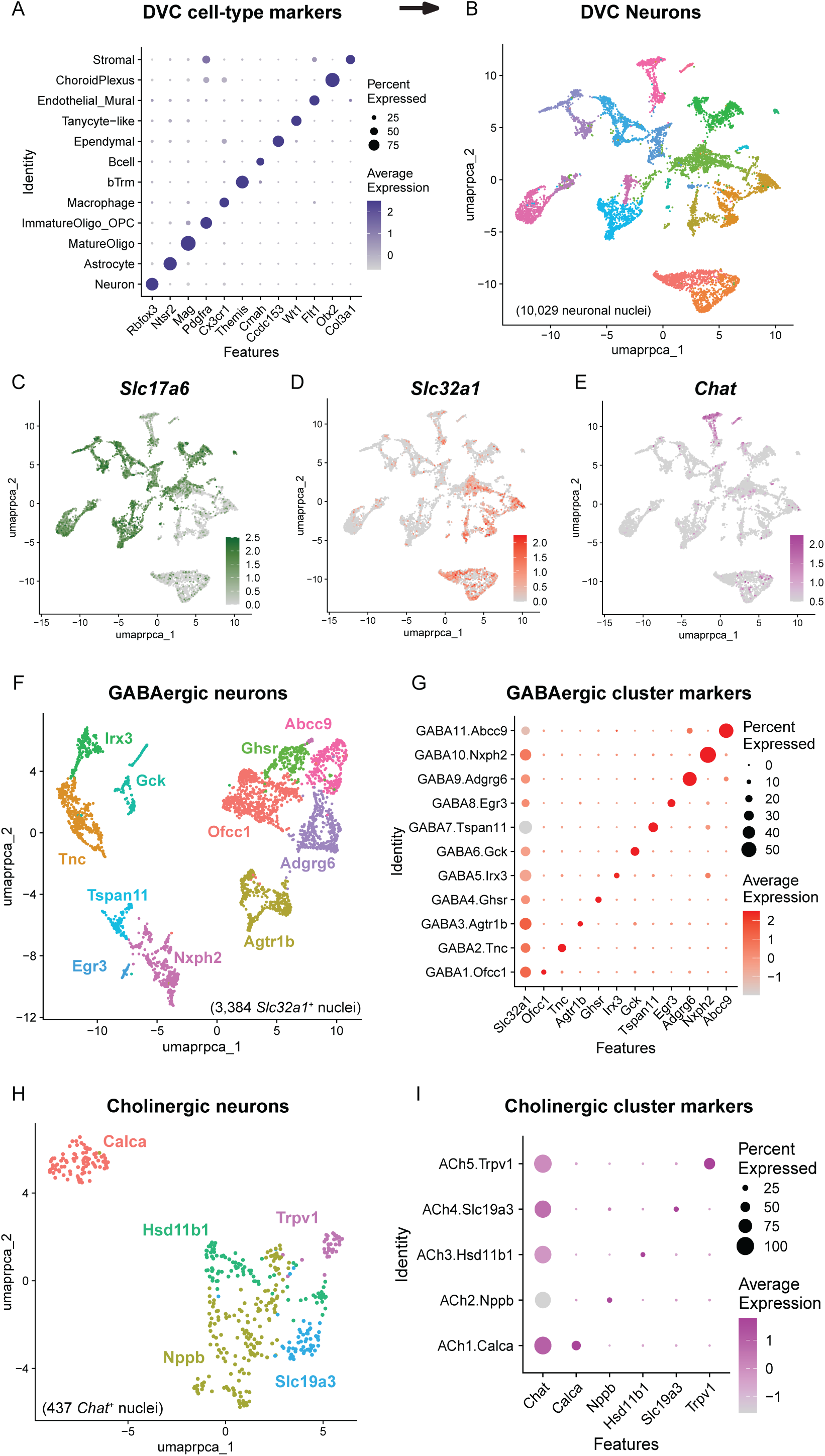
snRNA-seq profiling and marker gene analysis of dorsal vagal complex neurons, related to Figure 3. **A)** Dot plot showing DVC cell cluster markers. **B)** UMAP of DVC neurons, comprised of 10,029 nuclei. **C)** Feature plot depicting the expression of *Slc17a6* in DVC neurons. **D)** Feature plot depicting the expression of *Slc32a1* in DVC neurons. **E)** Feature plot depicting the expression of *Chat* in DVC neurons. **F)** UMAP of all GABAergic (*Slc32a1*^+^) neurons, comprising 3,384 nuclei. **G)** Dot plot showing *Slc32a1*^+^ cluster markers. **H)** UMAP of all cholinergic (*Chat*^+^) neurons, comprising 437 nuclei. **I)** Dot plot showing *Chat*^+^ cluster markers. DVC, dorsal vagal complex.

**Figure S4:**
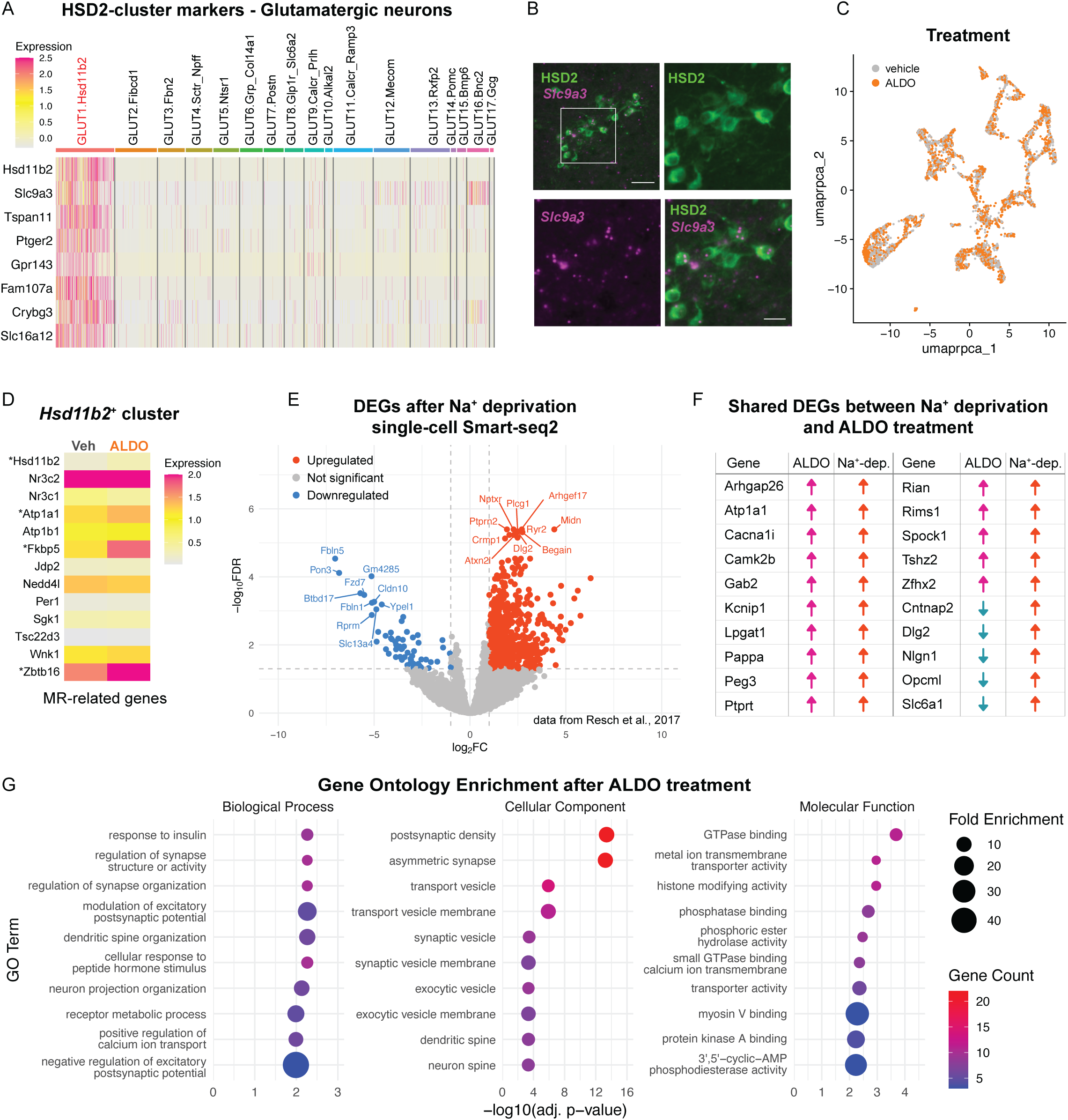
HSD2 neuron marker genes and differential gene expression induced by sodium deficiency and aldosterone, related to Figure 3. **A)** Heatmap depicting GLUT1.Hsd11b2 cluster markers in *Slc17a6*^+^ neurons. **B)** Top left panel shows low-magnification view of HSD2 immunofluorescence (IF) colocalized with *Slc9a3* fluorescence *in situ* hybridization (FISH; scale bar = 50 μm). Remaining panels provide high-magnification views of the boxed region for HSD2 IF (top right), *Slc9a3* FISH (bottom left), and both (bottom right; scale bar = 20 μm). **C)** UMAP of all *Slc17a6*^+^ neurons grouped by vehicle or aldosterone treatment. **D)** Heatmap depicting MR-related gene expression in the GLUT1.Hsd11b2 cluster after vehicle (left; Veh) or aldosterone (ALDO) treatment (right). * depicts statistically significant change in expression induced by ALDO. **E)** Volcano plot showing DEGs in scRNA-seq of HSD2 neurons following dietary sodium deprivation^14^. Downregulated genes are shown in blue and upregulated genes are shown in orange. In this Smart-seq2 study, genes with an absolute log_2_ fold change (log_2_FC) > 1 and adjusted false discovery rate (FDR) of < 0.05 were considered significant. **F)** Shared DEGs in HSD2 neurons between dietary sodium deprivation and aldosterone treatment. **G)** Bubble charts depicting top hits identified by Gene Ontology (GO) enrichment analysis of upregulated genes following aldosterone treatment. Benjamini-Hochberg adjustment was used to correct for multiple comparisons, and the top 10 GO terms with the lowest adjusted p-value are plotted for each GO category.

**Figure S5:**
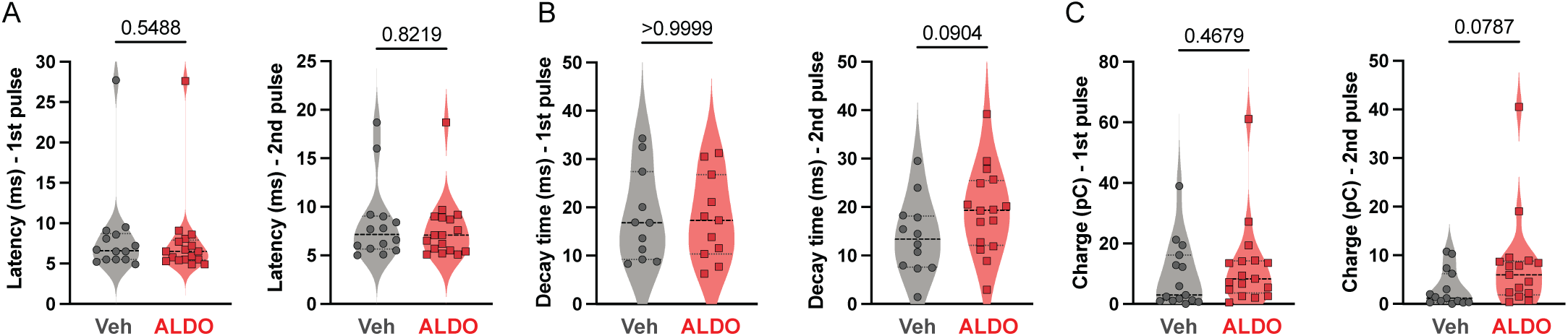
HSD2 neuron à BSTvL paired-pulse recording parameters and Sv2c expression in HSD2 neurons after CRISPR-Cas9 *Nr3c2* disruption, related to Figure 5. **A)** Light-evoked EPSC response latency to the first (left) and second (right) pulse (n = 14-17 neurons from 2-4 mice/group). **B)** Light-evoked EPSC response decay time after the first (left) and second (right) pulse (n = 11-16 neurons from 2-4 mice/group). **C)** Light-evoked EPSC charge from the first (left) and second (right) pulse (n = 15-17 neurons from 2-4 mice/group). All data are represented as median and quartiles. All statistical analyses were performed using a Welch’s t-test. p < 0.05 was considered statistically significant.

**Figure S6:**
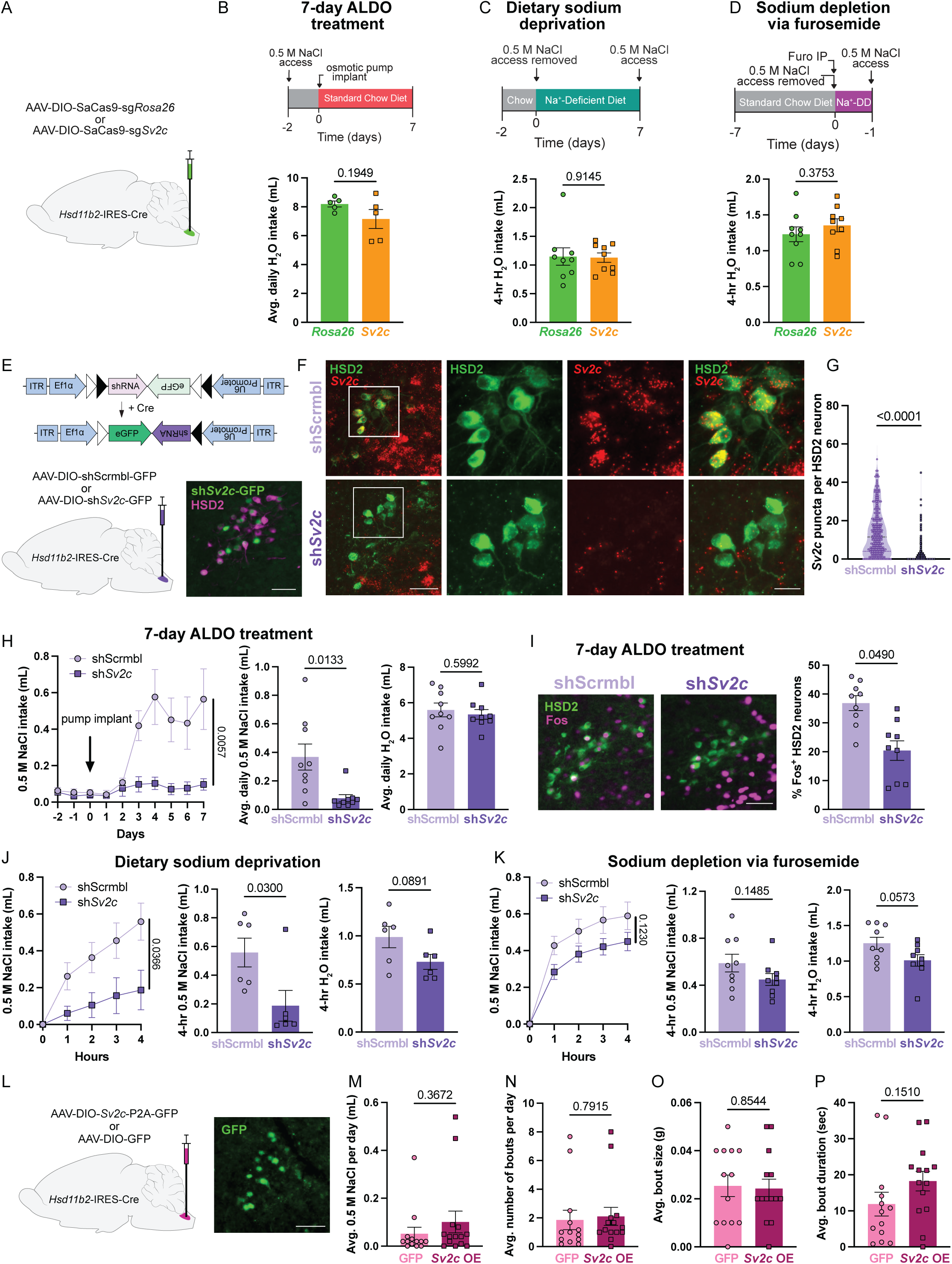
Saline and water intake after manipulation of *Sv2c* expression in HSD2 neurons, related to Figure 5. **A)** Schematic showing injection of Cre-dependent AAV-Cas9 targeting either *Rosa26* or *Sv2c* into *Hsd11b2*-IRES-Cre mice. **B)** Schematic describing the aldosterone-induced sodium appetite model (top) and average H_2_O intake per day (bottom; n = 5 mice/group). ALDO = aldosterone. **C)** Schematic describing the dietary sodium deprivation sodium appetite model with Na^+^-deficient diet (top) and total H_2_O intake in 4 hours (bottom; n = 9 mice/group). **D)** Schematic for the furosemide-induced sodium depletion sodium appetite model (top) and total H_2_O in 4 hours for each mouse (bottom: n = 9 mice/group). Furo = furosemide; Na^+^-DD = Na^+^-deficient diet. **E)** AAV diagram for Cre-dependent shRNA targeting *Sv2c* (top), schematic for Cre-dependent AAV-sh*Sv2c*-eGFP or AAV-shScramble-eGFP injection into *Hsd11b2*-IRES-Cre mice (bottom left), and representative histology of HSD2 and GFP immunofluorescence (IF; scale bar = 50 μm). **F)** Representative brain sections from HSD2^shScrmbl^ (top) or HSD2^sh*Sv2c*^ (bottom) mice following 7 days of aldosterone treatment showing, left to right: low-magnification view of HSD2 IF and *Sv2c* fluorescence *in situ* hybridization (FISH; scale bar = 50 μm); high-magnification of boxed region with HSD2 IF, high-magnification of boxed region with *Sv2c* FISH, and both HSD2 IF and *Sv2c* FISH (scale bar = 20 μm). **G)** Total number of *Sv2c* puncta per HSD2 neuron (n = 288-289 neurons from 3 mice/group). Data are represented as median and quartiles. **H)** 0.5 M NaCl intake in HSD2^shScrmbl^ or HSD2^sh*Sv2c*^ mice during 7 days of aldosterone treatment (left) and average 0.5 M NaCl (middle) and H_2_O (right) intake per day (n = 9 mice/group). **I)** Representative images showing c-Fos expression in HSD2 neurons of HSD2^shScrmbl^ (left) and HSD2^sh*Sv2c*^ (middle) mice following 7 days of aldosterone treatment and the percentage of HSD2 neurons expressing c-Fos (right; n = 9 mice/group; scale bar = 50 μm). Data shown as mean ± SEM. **J)** 0.5 M NaCl intake in HSD2^shScrmbl^ or HSD2^sh*Sv2c*^ mice over 4 hours following dietary sodium deprivation (left) and total 0.5 M NaCl (middle) and H_2_O (right) intake in 4 hours (n = 6 mice/group). **K)** 0.5 M NaCl intake in HSD2^shScrmbl^ or HSD2^sh*Sv2c*^ mice over 4 hours following sodium depletion via furosemide (left) and total 0.5 M NaCl (middle) and H_2_O (right) intake in 4 hours (n = 6 mice/group). **L)** Schematic showing injection of Cre-dependent AAV-SV2C-GFP or AAV-GFP into the NTS of *Hsd11b2*-IRES-Cre mice (left) and representative GFP IF indicating SV2C overexpression (right; scale bar = 50 µm). **M)** Average daily 0.5 M NaCl intake in HSD2^SV2C-OE^ or HSD2^GFP^ mice (n = 13-14 mice/group). **N)** Average number of bouts per day (n = 13-14 mice/group). **O)** Average bout size (n = 13-14 mice/group). **P)** Average bout duration (n = 13-14 mice/group). All 0.5 M NaCl and H_2_O intake data are represented as mean ± SEM. Statistical analysis of saline intake over time between groups was performed using a two-way repeated-measures ANOVA with Geisser-Greenhouse correction. All other statistical analyses were performed with a Welch’s t-test. p < 0.05 was considered statistically significant.

